# Inferring synaptic inputs from spikes with a conductance-based neural encoding model

**DOI:** 10.1101/281089

**Authors:** Kenneth W. Latimer, Fred Rieke, Jonathan W. Pillow

## Abstract

**A popular approach to the study of information processing in the nervous system is to characterize neural responses in terms of a cascade of linear and nonlinear stages: a linear filter to describe the neuron’s stimulus integration properties, followed by a rectifying nonlinearity to convert filter output to spike rate. However, real neurons integrate stimuli via the modulation of nonlinear excitatory and inhibitory synaptic conductances. Here we introduce a bio-physically inspired point process model with conductance-based inputs. The model provides a novel interpretation of the popular Poisson generalized linear model (GLM) as a special kind of conductance-based model, where excitatory and inhibitory conductances are modulated in a “push-pull” manner so that total conductance remains constant. We relax this constraint to obtain a more general and flexible “conductance-based encoding model” (CBEM), which can exhibit stimulus-dependent fluctuations in gain and dynamics. We fit the model to spike trains of macaque retinal ganglion cells and show that, remarkably, we can accurately infer underlying inhibitory and excitatory conductances, using comparisons to intracellularly measured conductances. Using extracellular data, we corroborate the intracellular finding that synaptic excitation temporally precedes inhibition in retina. We show that the CBEM outperforms the classic GLM at predicting retinal ganglion cell responses to full-field stimuli, generalizes better across contrast levels, and captures inhibition-dependent response properties to spatially structured stimuli. The CBEM provides a powerful tool for gaining insights into the intracellular variables governing spiking, and forges an important link between extracellular characterization methods and biophysically detailed response models.**

## 1 Introduction

Studies of neural coding in the early sensory pathway seek to reveal how sensory information is encoded in neural responses. A complete understanding this code requires knowledge of the statistical relationship between stimuli and spike trains as well as the biophysical mechanisms by which this transformation is carried out. A popular approach to the neural coding problem is to use “cascade” models such as the linear-nonlinear-Poisson (LNP) or generalized linear model (GLM) to characterize how external stimuli are converted to spike trains. These descriptive statistical models describe the encoding process in terms of a series of stages: linear filtering, nonlinear transformation, and noisy spiking (Chichilnisky, 2001; Paninski, 2004; Butts et al., 2011; Vintch et al., 2012; McFarland et al., 2013; Park et al., 2013a; Theis et al., 2013; Vintch et al., 2015). The Poisson GLM in particular has provided a powerful tool for characterizing neural encoding in a variety of sensory, cognitive, and motor brain areas (Harris et al., 2003; Truccolo et al., 2005; Pillow et al., 2008; Gerwinn et al., 2010; Stevenson et al., 2012; Weber et al., 2012; Park et al., 2014; Hardcastle et al., 2015; Yates et al., 2017).

However, there is a substantial gap between descriptive statistical models like the GLM and mechanistic or biophysically interpretable models. In real neurons, stimulus integration is nonlinear, arising from excitatory and inhibitory synaptic inputs that depend nonlinearly on the stimulus. These synaptic inputs drive conductance changes that alter the nonlinear dynamics governing membrane potential. In retina and other sensory areas, the tuning of excitatory and inhibitory inputs can differ substantially (Roska et al., 2006; Trong & Rieke, 2008; Poo & Isaacson, 2009; Cafaro & Rieke, 2013). Determining how these inputs influence the spiking outputs of neurons, and thus the computations performed by the cells, remains an important challenge. This challenge is of course exacerbated by the fact that most studies of neural coding rely on extracellular recording techniques, which detect only spiking activity and not synaptic conductances.

Here we aim to narrow the gap between descriptive statistical and biophysically interpretable models, while remaining within the domain of models that can be estimated from extracellular spike train data (Meng et al., 2011, 2014; Lankarany, 2017). We first introduce a quasi-biophysical interpretation of the standard Poisson GLM, which reveals its equivalence to a constrained conductance-based model with equal and opposite excitatory and inhibitory tuning. We then expand on this interpretation to formulate a more flexible and realistic conductance-based model with independent tuning of excitatory and inhibitory inputs, which we refer to as the conductance-based encoding model. This model can mimic real neurons in exhibiting shunting as well as hyperpolarizing inhibition, and time-varying changes in gain and membrane time constant.

To validate the CBEM’s biophysical and statistical accuracy, we applied it to data recorded in macaque retinal ganglion cells. We show, using intracellular data, that the stimulus-driven excitatory and inhibitory conductances in these neurons were well characterized by linear-nonlinear cascades, with the filter driving inhibition exhibiting a slight delay relative to the filter driving excitation. Moreover, the CBEM allowed for accurate inference of excitatory and inhibitory synaptic conductances from spike trains alone, using comparisons to intracellular conductance measured from the same same cell. Finally, we show that the CBEM predicted retinal spike responses more accurately than the standard GLM for both full-field and spatio-temporally varying stimuli.

## 2 Background: Poisson GLM with spike history

Generalized linear models provide a simple yet powerful description of the encoding relationship between stimuli and neural responses (Truccolo et al., 2005). A recurrent Poisson generalized linear model, often referred to in the neural coding literature simply as the “GLM”, describes neural encoding in terms of a cascade of linear, nonlinear, and probabilistic spiking stages (***Fig. 1a***). The GLM parameters include a stimulus filter **k**, which describes how the neuron integrates an external stimulus, a post-spike filter **h**, which captures dependencies on spike history, and a baseline *b* that determines baseline firing rate in the absence of input. The outputs of these filters are summed and passed through a nonlinear function *f*_*r*_ to obtain the conditional intensity for an inhomogeneous Poisson spiking process. The model can be written concisely in discrete time as:

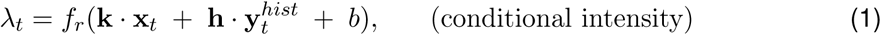

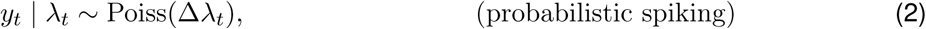

where **x**_*t*_ is the spatio-temporal stimulus vector at time *t*, 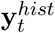 is a vector of spike history at time *t, λ*_*t*_ *≥* 0 is the spike rate at time *t*, and *y*_*t*_ is the spike count in bin of size Δ. Although spike generation is conditionally Poisson, the model can capture complex history-dependent response properties such as refractoriness, bursting, bistability, and adaptation (Weber & Pillow, 2016). Additional filters can be added to the model in order to incorporate dependencies on covariates of the response such as spiking in other neurons or local field potential recorded on nearby electrodes (Truccolo et al., 2005; Pillow et al., 2008; Kelly et al., 2010). A common choice for the nonlinearity is exponential, *f* (*z*) = exp(*z*), which corresponds to the “canonical” inverse link function for Poisson GLMs.

**Figure 1:**
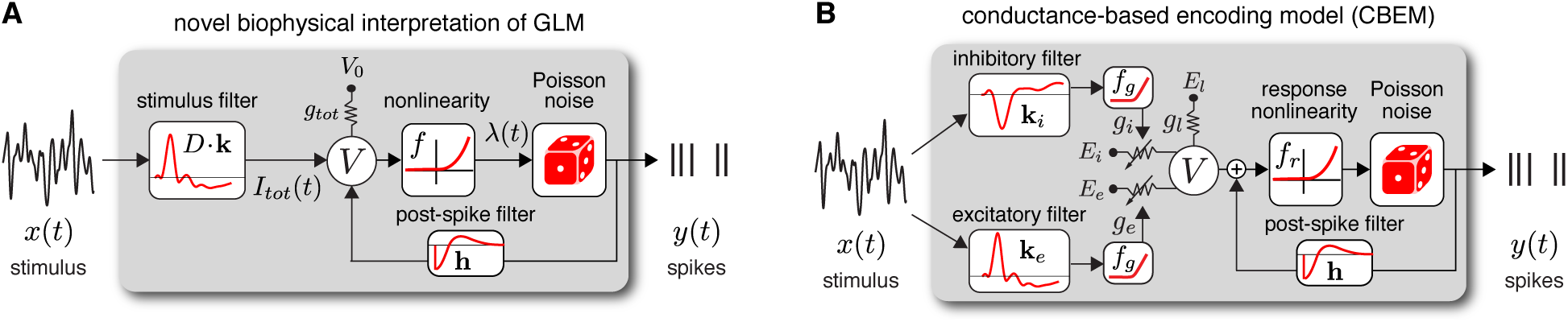
Model diagrams. **(A)** Diagram illustrating novel biophysical interpretation of the generalized linear model (GLM). The stimulus *x*_*t*_ is convolved with a conductance filter **k** weighted by *D* = (*E*_*e*_ *− E*_*i*_), the difference between excitatory and inhibitory current reversal potentials, resulting in total synaptic current *I*_*tot*_(*t*). This current is injected into the linear RC circuit governing the membrane potential *V*_*t*_, which is subject to a leak current with conductance *g*_*tot*_ and reversal potential *V*>_0_. The instantaneous probability of spiking is governed by a the conditional intensity *λ*_*t*_ = *f* (*V*_*t*_), where *f* is a nonlinear function with non-negative output. Spiking is conditionally Poisson with rate *λ*_*t*_, and spikes gives rise to a post-spike current or filter **h** that affects the subsequent membrane potential. **(B)** Conductance-based encoding model (CBEM). The stimulus **x**_*t*_ is convolved with filters **k**_*e*_ and **k**_*i*_, whose outputs are transformed by rectifying nonlinearity *f*_*g*_ to produce excitatory and inhibitory synaptic conductances *g*_*e*_(*t*) and *g*_*i*_(*t*). These time-varying conductances and the static leak conductance *g*_*l*_ drive synaptic currents with reversal potentials *E*_*e*_, *E*_*i*_, and *E*_*l*_, respectively. The resulting membrane potential *V*_*t*_ is added to a linear spike-history term, given by **h** · **y**^*hist*^(*t*), and then transformed via rectifying nonlinearity *f*_*r*_ to obtain the conditional intensity *λ*_*t*_, which governs conditionally Poisson spiking as in the GLM.

Previous literature has offered a quasi-biological interpretation of the GLM known as “soft threshold” integrate-and-fire (IF) model (Plesser & Gerstner, 2000; Gerstner, 2001; Paninski et al., 2007; Mensi et al., 2011). This interpretation views the summed filter outputs as the neuron’s membrane potential, similar to the standard IF model, where membrane potential is a linearly fitered version of input cur-rent. The nonlinear function *f*_*r*_ can be interpreted as a “soft threshold” function that governs a smooth increase in the instantaneous spike probability as a function of membrane depolarization. Lastly, the post-spike current **h** determines how membrane potential is reset following a spike.

We can rewrite the standard GLM to emphasize this biological interpretation explicitly:

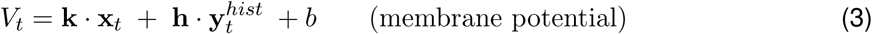

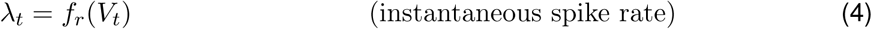

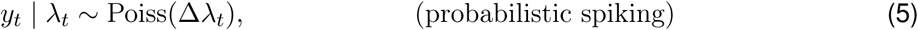

Note that in this “soft” version of the IF model, the only noise source is the conditionally Poisson spiking mechanism; this differs from other noisy extensions of the IF model with “hard” spike thresholds that require more elaborate methods for computing likelihoods (Paninski, 2004; Pillow et al., 2005; Paninski et al., 2008). To convert this model to a classic leaky integrate-and-fire model, we could replace *f*_*r*_ with a “hard” threshold function that jumps from zero to infinity at some threshold value of the membrane potential, set the stimulus filter **k** to an exponential decay filter, and set the post-spike filter **h** to a delta function that causes instantaneous reset of the membrane potential following a spike. The GLM membrane potential is a linear function of the input, just as in the classic leaky IF model, and thus both models fail to capture the nonlinearities apparent in the synaptic inputs to most real neurons (Schwartz & Rieke, 2011).

## 3 Interpreting the GLM as a conductance-based model

Here we propose a more biophysically realistic interpretation of the GLM by moving to a dynamical model with conductance-based input. Consider a neuron with membrane potential *V*_*t*_ governed by the ordinary differential equation (ODE):

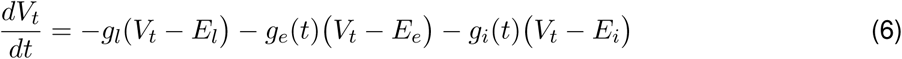

where *g*_*l*_ is leak conductance, *g*_*e*_(*t*) and *g*_*i*_(*t*) are time-varying excitatory and inhibitory synaptic conductances, and *E*_*l*_, *E*_*e*_ and *E*_*i*_ are the leak, excitatory and inhibitory reversal potentials. Note that we have ignored capacitance, which would provide an (unobserved) scaling factor on *dV/dt*, but will not affect our results. We can rewrite this model in terms of a “total conductance” given by

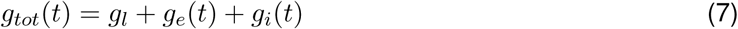

and an effective total input current given by

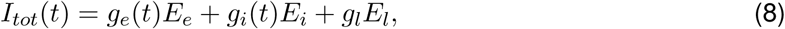

simplifying the membrane equation to:

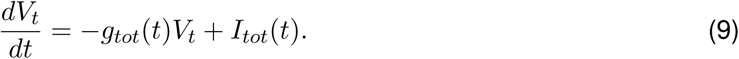

A natural question to ask is: under what conditions, if any, is this model a GLM? Here we provide a set of sufficient conditions for an equivalence between the two. The definition of a GLM requires membrane potential *V*_*t*_ to be a linear function of the stimulus, which holds if the two following conditions are met:

1. Total conductance *g*_*tot*_(*t*) is constant, making (***Eq. 9***) a linear ODE.
2. The input current *I*_*tot*_(*t*) is an affine (linear plus constant) function of the stimulus **x**_*t*_.

The first condition implies *g*_*e*_(*t*) + *g*_*i*_(*t*) = *c*, for some constant *c*, and the second implies that *g*_*e*_(*t*)*E*_*e*_ + *g*_*i*_(*t*)*E*_*i*_ is a linear function of the stimulus. We can satisfy these two conditions simultaneously by modeling the excitatory and inhibitory conductances as linear (or affine) functions of the stimulus with identical filters of opposite sign:

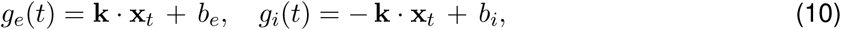

where *b*_*e*_ and *b*_*i*_ are arbitrary constants. Under this setting, excitatory and inhibitory conductances are driven by equal and opposite linear projections of the stimulus, with total conductance fixed at *g*_*tot*_ = *g*_*l*_ + *b*_*e*_ + *b*_*i*_ and synaptic input equal to *I*_*tot*_(*t*) = (*E*_*e*_ *− E*_*i*_)(**k** · **x**_*t*_) + *E*_*e*_*b*_*e*_ + *E*_*i*_*b*_*i*_. We then rewrite the membrane equation (***Eq. 6***) as:

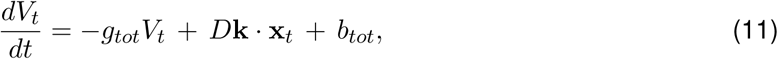

where *g*_*tot*_ = *g*_*l*_ + *b*_*e*_ + *b*_*i*_ is the (constant) total conductance, *D* = *E*_*e*_ *− E*_*i*_ scales the linear filter output, and *b*_*tot*_ = *b*_*e*_*E*_*e*_ + *b*_*i*_*E*_*i*_. If the initial voltage is set to *V*_0_ = *b*_*tot*_*/g*_*tot*_, then the membrane potential is given by

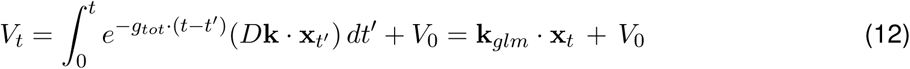

where the equivalent standard GLM filter is equal to the linear convolution of **k** with an exponential decay filter **k**_*leak*_:

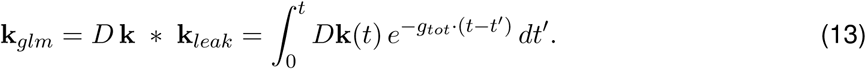

The membrane potential *V*_*t*_ is a linear function of the stimulus, so if we include a monotonic nonlinearity and conditionally Poisson spiking, this model is clearly a GLM.

Thus, to summarize: the GLM can be interpreted as a conductance-based dynamical model in which a common linear “conductance” filter drives equal and opposite fluctuations in excitatory and inhibitory synaptic conductances; the GLM filter **k**_*glm*_ is equal to the convolution of this conductance filter with an exponential decay filter whose time constant is the inverse of the total conductance.

## 4 The conductance-based encoding model (CBEM)

From this novel interpretation of the GLM, it is straightforward to formulate a more realistic conductance-based statistical spike train model. Namely, we can remove the constraint that excitatory and inhibitory conductance sum to a constant. Relaxing this constraint, so that total conductance van vary, results in a new model that we refer to as the *conductance-based encoding model* (CBEM). The CBEM represents an extension of GLM to allow for differential tuning of excitation and inhibition (i.e., **k**_*e*_ ≠ −**k**_*i*_), and adds rectifying nonlinearities governing the relationship between the stimulus and synaptic conductances. (See model diagram, ***Fig. 1b***). The CBEM model is no longer a GLM because the filtering it performs on the stimulus is nonlinear.

Formally, the CBEM is driven by excitatory and inhibitory synaptic conductances that are each linear-nonlinear functions of the stimulus:

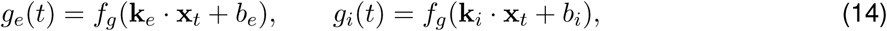

where **k**_*e*_ and **k**_*i*_ are linear filters driving excitatory and inhibitory conductance, respectively, *f*_*g*_ is a rec-tifying nonlinearity to ensure that conductances are non-negative, and *b*_*e*_ and *b*_*i*_ determine the baseline excitatory and inhibitory conductances in the absence of input. We will assume a “soft-rectification” nonlinearity given by

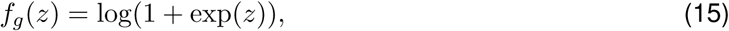

which behaves like a smooth version of a linear half-rectification function.

The CBEM membrane potential *V*_*t*_ then evolves according to the nonlinear ordinary differential equation (***Eq. 6***) under the influence of the two time-varying conductances *g*_*e*_(*t*) and *g*_*i*_(*t*). We use a first-order exponential integrator method to solve this equation, which is exact under the assumption that *g*_*e*_(*t*) and *g*_*i*_(*t*) are constant during each time bin of size Δ, giving updates:

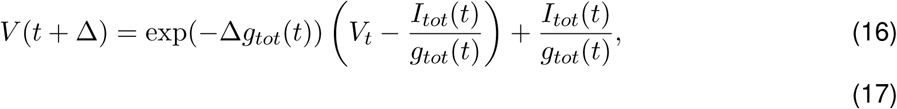

for *g*_*tot*_(*t*) and *I*_*tot*_(*t*) in ***Eqs. 7-8*** and assuming *V* (0) = *E*_*l*_ at the start of each experiment.

To incorporate spike-history dependencies, we add a linear autoregressive term to obtain an “effective” membrane potential

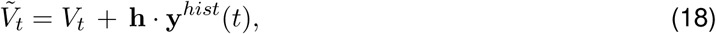

where **y**^*hist*^(*t*) is a vector of previous spike history at time *t*. Although one could add this autoregressive spike-history term to the membrane potential equation (***Eq. 6***), this implementation allowed us to focus our analyses on stimulus-induced as opposed to spike-induced changes in conductance.

The effective membrane potential 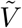 is transformed to instantaneous spike rate *λ*(*t*) by an output nonlinearity *f*_*r*_(·). We selected a biophysically motivated output nonlinearity proposed by Mensi et al. (2011):

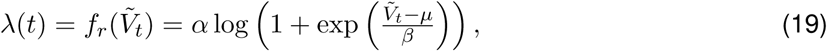

where *µ* is a “soft” spike threshold, and *α* and *β* jointly determine slope and sharpness of the nonlinearity, respectively. Spiking is a conditionally Poisson process with conditional intensity *λ*(*t*).

The CBEM is similar to the Poisson GLM in that the only source of stochasticity is the conditionally Poisson spiking mechanism: we assume no additional noise in the conductances or the voltage. This simplifying assumption, although not biophysically accurate, makes log-likelihood simple to compute, allowing for efficient maximum likelihood inference using standard ascent methods (See Methods).

## 5 Validating the model with intracellular recordings

Before applying the CBEM to extracellularly recorded spike trains, we sought to validate our modeling assumptions using experiments in which spikes and conductances were measured from the same cells.

### 5.1 Conductance model

The CBEM describes the excitatory and inhibitory synaptic input received by a neuron in terms of a linear function of the stimulus followed by a point nonlinearity. A priori, this linear-nonlinear assumption is plausible for retinal ganglion cells (RGCs) because these cells receive excitatory synaptic input from bipolar cells, which exhibit membrane potential responses to full-field visual stimuli that are well-characterized by linear-nonlinear models (Rieke, 2001; Demb et al., 2001; Beaudoin et al., 2008; Gollisch & Meister, 2010). We therefore analyzed voltage clamp recordings from ON parasol RGCs in response to a full-field noise stimulus (Trong & Rieke, 2008). Active conductances intrinsic to the RGC were blocked during these recordings and the holding potential was set to isolate either the excitatory or inhibitory inputs received by the cell.

We fit the measured conductances with a linear-nonlinear cascade model (***Fig. 2***). We fixed the nonlinearity to be a soft-rectified nonlinearity (***Eq. 15***) which provided a close approximation to the relationship between projected stimulus and observed conductances (Liu et al., 2017; Real et al., 2017). We found these LN models accounted for 79 ± 4% (mean ± SEM) of the variance of the average excitatory conductance and 63 ± 3% of the inhibitory conductances recorded in response to a novel stimulus.

**Figure 2:**
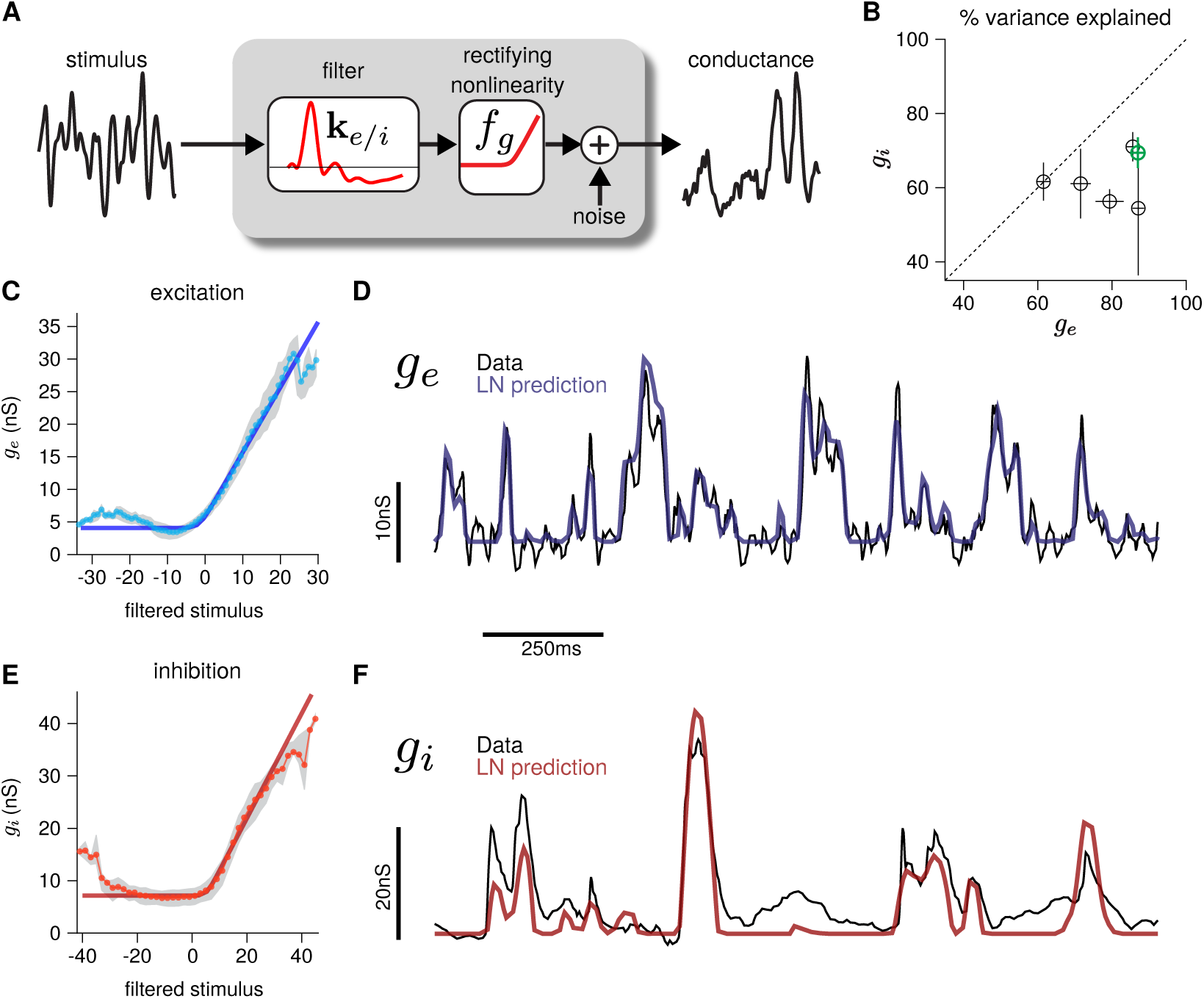
Validating the LN conductance model. The CBEM describes the relationship between stimulus and each synaptic conductance with a linear-nonlinear (LN) cascade, consisting of a linear filter followed by fixed rectifying nonlinearity. To validate this assumption, we fit LN models directly to measured excitatory and inhibitory conductances from a parasol retinal ganglion cell (RGC), fixing the nonlinearity to a soft-rectification function (***Eq. 15***) and optimizing the filter to minimize the mean squared error. (**A**) LN conductance model diagram. (**B**) The percent variance explained (*R*^2^) for excitatory and inhibitory conductances from 6 ON parasol RGCs, computed using cross-validation with a 6s test stimulus. Error bars indicate standard deviation of the variance explained across all test stimuli. (**C**) The excitatory conductance as a function of the filtered stimulus values for the example cell indicated in green in B. The gray region shows the middle 50-percentile of the distribution of observed excitatory conductance given the filtered stimulus value. The fixed soft-rectifying function (dark blue) closely matched the average conductance given the filtered stimulus value (light blue points). (**D**) Measured excitatory conductances (black) are compared to the predictions from the LN model (blue) on a withheld test stimulus for the cell shown in C. (**E**) The inhibitory conductances conditioned on the filtered stimulus value for the same example cell as in C. The fixed soft-rectifying function (dark red) provided a close approximation to the average inhibitory conductance given the filtered stimulus value (light red points). (**F**) Measured excitatory conductances (black) are compared to the predictions from the LN model (red) on a withheld test stimulus for the same cell.

Although this simplified model does not account for all the features of the synaptic conductances in RGCs (e.g., nonlinear spatial integration (Schwartz et al., 2012; Turner & Rieke, 2016), presynaptic gain control (Beaudoin et al., 2008; Cui et al., 2016a)), it accurately captures the dependence of synaptic input on a stationary full-field noise stimulus (Butts et al., 2016). Moreover, its simplicity enables computationally efficient fitting of the CBEM to extracellularly recorded spike trains, which we discuss below.

### 5.2 Spike rate nonlinearity

The second component of the CBEM is a nonlinearity, *f*_*r*_, mapping intracellularly recorded membrane voltage to instantaneous spike rate. To validate this modeling assumption, we examined dynamic current clamp recordings from two ON parasol RGCs (***Fig. 3a***). The dynamic clamp recordings drove RGCs with currents determined by previously measured excitatory and inhibitory conductances. To reduce noise, we computed average membrane potential over repeated presentations of the same measured conductance traces. We then computed nonparametric estimates of the nonlinearity for each conductance by taking a ratio of histograms: we used histograms to estimate the conditional distribution of the membrane potential given a spike, *P* (*V* |spike), and the marginal distribution over membrane potential, *P* (*V*), binned in 0.5 mV intervals (***Fig. 3b***), and then applied Bayes’ rule to obtain the nonlinearity as *P* (spike*|V*) = *P* (*V* |spike)*P* (spike)*/P* (*V*), where *P* (spike) is the number of spikes divided by the number of time bins. To obtain the nonlinearity in units of spikes/s, we divided by the time bin size (0.1 ms) (***Fig. 3c***). This is the same procedure for estimating the LN-model nonlinearity proposed in (Chichilnisky, 2001; Mease et al., 2013), but substituting the filtered stimulus with the average voltage measured across trials. We found that the parametric nonlinearity we assumed (red, ***Eq. 19***) closely approximated the non-parametric estimate (black; see Methods for details).

**Figure 3:**
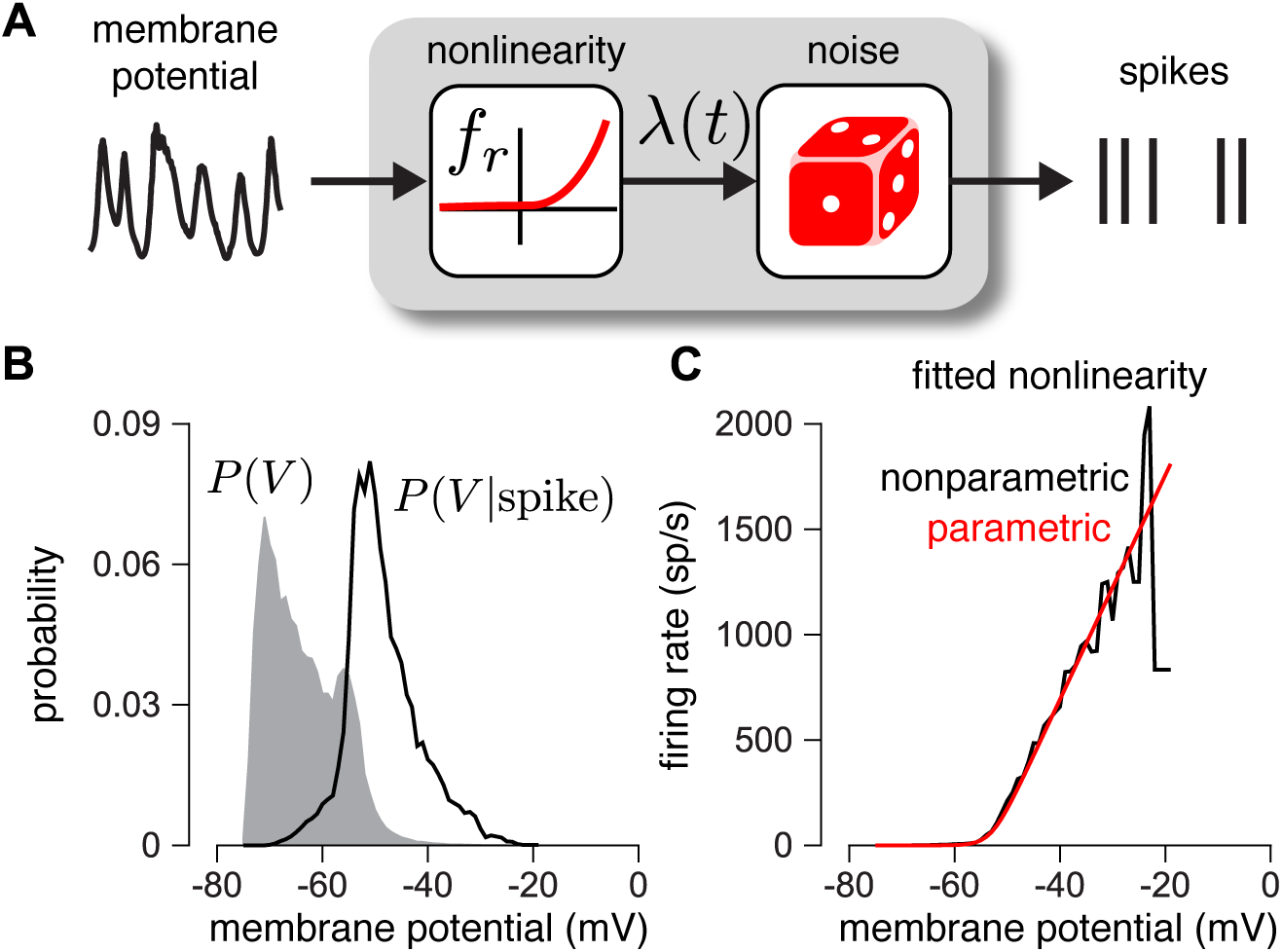
Validating the firing rate nonlinearity. (**A**) Schematic of the transformation from voltage to firing rate of a neuron. The mean voltage recorded during repeated trials of current injection is passed through a nonlinear function to predict the spike rate. (**B**) The distribution of mean membrane potential over several repeats of conditioned on a spike occurring during one of the repeats (black trace) for two parasol RGCs. The grey distribution shows the marginal distribution over membrane potential. (**C**) The firing rate as a function of voltage (black trace) computed by applying Bayes’ rule on the distributions shown in B. The firing rate function is closely approximated by a rectified linear function (red trace).

We note that previous analyses of RGC responses using Poisson GLMs have shown that an exponential nonlinearity captures the mapping from stimuli to spike rates more accurately than a rectified-linear nonlinearity (Pillow et al., 2008). Here we found the opposite: the nonlinearity was better described with a soft-rectification function. This discrepancy may result from the fact that the GLM has a single nonlinearity, whereas the CBEM has a cascade of two nonlinearities: one mapping filter output to conductance, and a second mapping membrane potential to spike rate.

## 6 Applying CBEM: inferring conductances from spikes

We now turn to the key application of the CBEM: inferring intracellular excitatory and inhibitory conductances from extracellular spike train data. To test our ability to make such predictions, we fit the CBEM parameters to a training dataset consisting of spike times elicited in response to a set of training stimuli, then used the inferred model filters to predict the excitatory and inhibitory conductances elicited in response to novel stimuli.

The training data consisted of spike trains from 6 macaque ON-parasol RGCs obtained in cell-attached recordings with full-field white noise stimuli. Each cell was stimulated with ten unique 6s stimulus segments, repeated 3 or 4 times each, resulting in a total of thirty to forty 6s trials per neuron (Trong & Rieke, 2008). We fit the CBEM parameters (conductance filters and spike history filter) responses to 9 of the stimulus segments and evaluated performance using the remaining held-out segment (10-fold cross validation). Thus, the model was fit using spike trains elicited by 3 or 4 repeats of a 54 second full-field noise stimulus (see Methods). For comparison, we also fit the CBEM conductance filters using measured excitatory and inhibitory conductances from intracellular recordings using the same stimuli.

***Fig. 4*** shows the conductance filters estimated from intracellular data (fit to conductances) and extracellular data (fit to spike trains only) for two example cells, along with the predicted excitatory and inhibitory conductances elicited by a novel test stimulus. The filters fit to spikes were similar to those fit to conductances, and the conductance predictions from both models were highly correlated with the measured traces. ***Fig. 5*** shows a summary statistics comparing the two models’ performance for all 6 neurons for which we had both spike train and conductance recordings. For both models, predicted conductances traces were highly correlated with the measured conductances for all 6 cells. Using only a few minutes of spiking data, the conductances predicted by the extracellular model had an average correlation of *r* = 0.73 ± 0.01 (mean ± SEM) for the excitatory conductance and *r* = 0.69 ± 0.03 for the inhibitory conductance, compared to averages of *r* = 0.89 ± 0.02 (excitation) and *r* = 0.82 ± 0.01 (inhibition) for the LN model fit directly to conductances (***Fig. 5a-b***).

**Figure 4:**
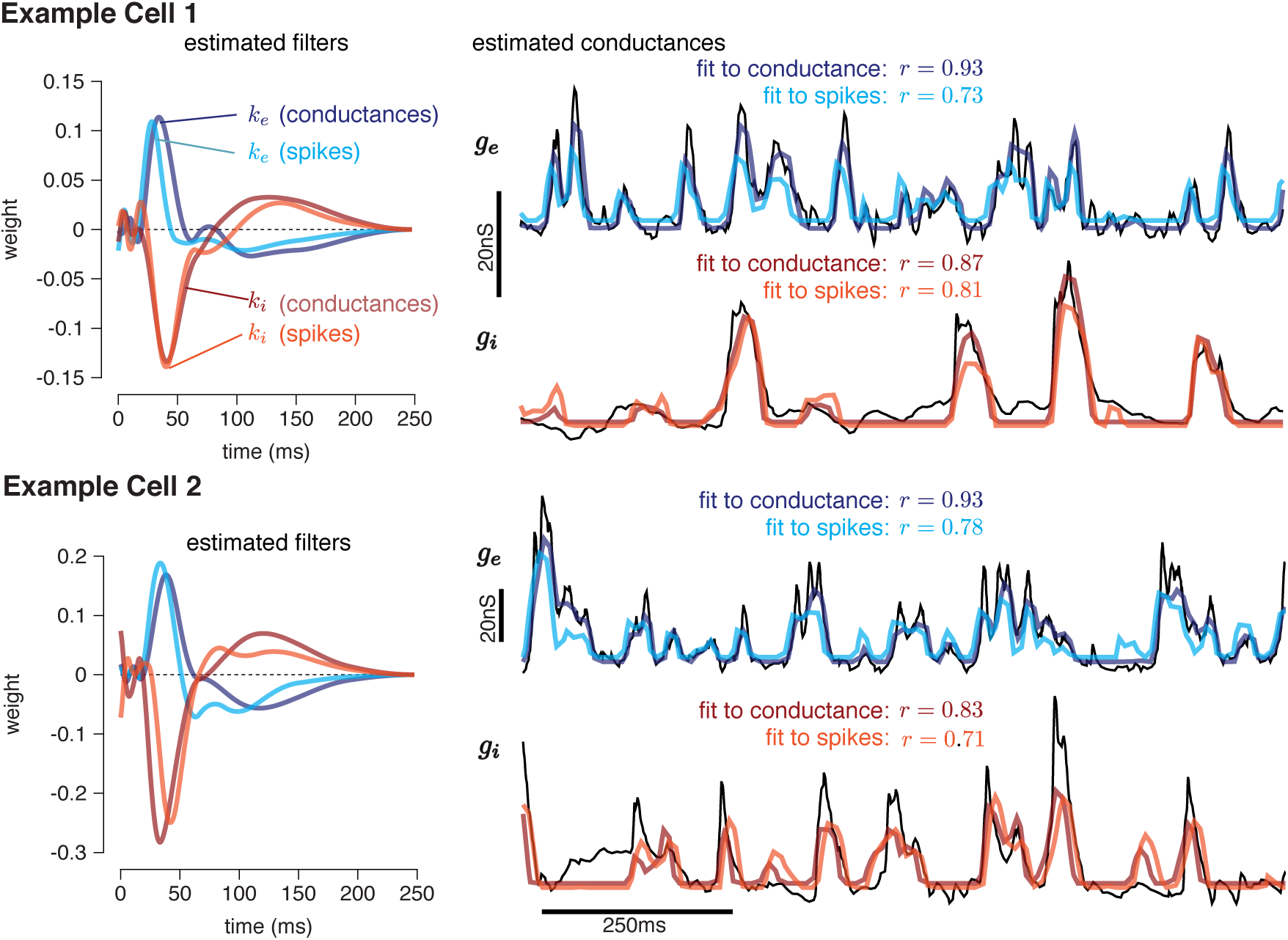
Two example ON parasol cells fit with the CBEM. We fit the CBEM parameters to spike train data and then used the inferred excitatory and inhibitory filters to predict excitatory and inhibitory synaptic currents recorded separately in response to novel stimuli. For comparison, we show predictions of an LN model fit directly to the conductance data. **Left**: Linear kernels for the excitatory (blue) and inhibitory (red) inputs estimated from the conductance-based model (light red, light blue) and estimated by fitting a linear-nonlinear model directly to the measured conductances (dark red, dark blue). The filters represent a combination of events that occur in the retinal circuitry in response to a visual stimulus, and are primarily shaped by the cone transduction process. **Right**: Conductances predicted by our model on a withheld test stimulus. Measured conductances (black) are compared to the predictions from the CBEM filters (fit to spiking data) and an LN model (fit to conductance data). To visualize the measured conductances alongside the inferred conductances, we scaled the estimated conductances. Real membrane voltage dynamics depend on the capacitance of the membrane, which we do not include because it introduces an arbitrary scaling factor that cannot be estimated from spikes alone. Therefore, for comparisons we chose a scaling factor for each cell to minimize the squared error between the predicted conductances and the measured conductances, while assuming a common scaling for the inhibitory and excitatory conductances. We similarly scaled the stimulus filters fit to the conductance measurements to match the average height of the CBEM filters.

**Figure 5:**
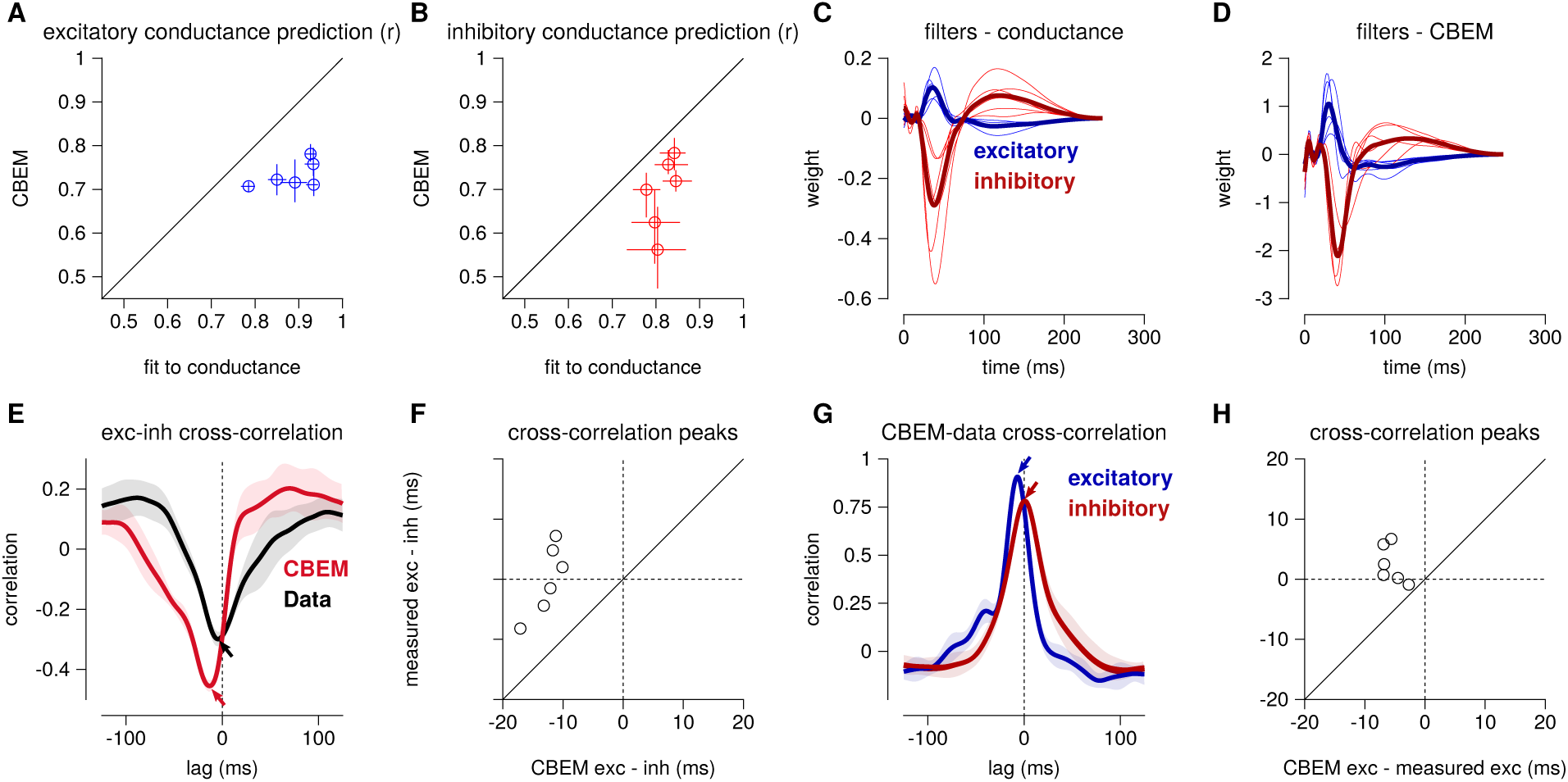
Summary of the CBEM fits to 6 ON parasol RGCs for which we had both spike train and conductance recordings. (**A**) The correlation coefficient (*r*) between the mean observed excitatory synaptic input to a novel 6 s stimulus and the conductance predicted by the LN cascade fit to the excitatory conductance (y-axis) compared to the CBEM prediction from spikes (x-axis) for each cell. Error bars indicate the minimum and maximum values observed across all cross-validated stimuli (**B**) Same as C for the inhibitory conductance. (**C**) The excitatory (blue) and inhibitory (red) filters estimated from voltage-clamp recordings. The thick traces show the mean filters. (**D**) The excitatory (blue) and inhibitory (red) filters estimated by the CBEM from spike trains. (**E**) The cross-correlation of the excitatory and inhibitory conductances for an example cell measured from the data (black trace; region shows standard deviation across the 10 stimuli) compared to the cross-correlation in the CBEM fit to that cell (red trace). Arrows indicate the peaks of the cross-correlations. In the data, excitation and inhibition are anti-correlated and show similar timing. However, excitation precedes inhibition in the model. (**F**) The cross-correlation peak times between excitation and inhibition measured from data (y-axis) compared to the conductances predicted by the CBEM (x-axis) for all 6 cells. Negative values on the x-axis indicate that excitation leads inhibition in the CBEM fits to these cells. (**G**) Comparing the timing of excitatory and inhibitory conductances from the data and the CBEM for the example cell in E. The cross-correlation between the measured excitatory conductance and the CBEM’s excitatory conductance (blue) and the cross-correlation between data and model for the inhibitory conductances (red). (**H**) Cross-correlation peak times between measured and CBEM predicted inhibition (y-axis) and excitation (x-axis).

Although the extracellular model predicted the basic timecourse of the observed conductances with high fidelity, there were small systematic discrepancies between model-predicted and measured con-ductances. For example, measured conductances had nearly zero lag in their cross-correlation (0.0 ± 2.4 ms; see also Cafaro & Rieke, 2013), whereas the predicted excitatory conductance slightly preceded the inferred inhibition for all 6 cells (12.6 ± 1.0 ms, Student’s *t*–test *p <* 0.0001; ***Fig. 5e-f***). The predicted excitation preceded the average measured excitation by 5.6 ± 0.7 ms (*p* = 0.0005), while the predicted inhibition showed only a slight and statistically insignificant delay compared to the measured inhibition (2.5 ± 1.3 ms, *p* = 0.11; ***Fig. 5g-h***).

## Characterizing spike responses with CBEM

Given the CBEM’s ability to infer intracellular conductances from spike train data, we sought to examine how well it predicts spike responses. Most encoding models are only ever fit and tested using extracellular recordings, which are far easier to perform and to sustain over longer periods. It therefore seems natural to ask: does the CBEM’s increased degree of biophysical realism confer advantages for predicting spikes?

To answer this question, we fit the CBEM and classic Poisson GLM to a population of 9 extracellularly recorded macaque RGCs stimulated with full-field binary white noise (Uzzell & Chichilnisky, 2004; Pillow et al., 2005). We evaluated spike rate prediction by comparing the peri-stimulus time histogram (PSTH) of the simulated models to the PSTH of real neurons using a 5s test stimulus (***Fig. 6)***. The CBEM had higher prediction accuracy than the GLM for all nine cells, capturing 86% of the variance of the PSTH on average vs. 77% for the GLM. We then evaluated spike train prediction by comparing log-likelihood on a 5 minute test dataset. The CBEM again outperformed the GLM on all cells, offering an improvement of 0.34 ± 0.11 bits/spike on average over the GLM.

**Figure 6:**
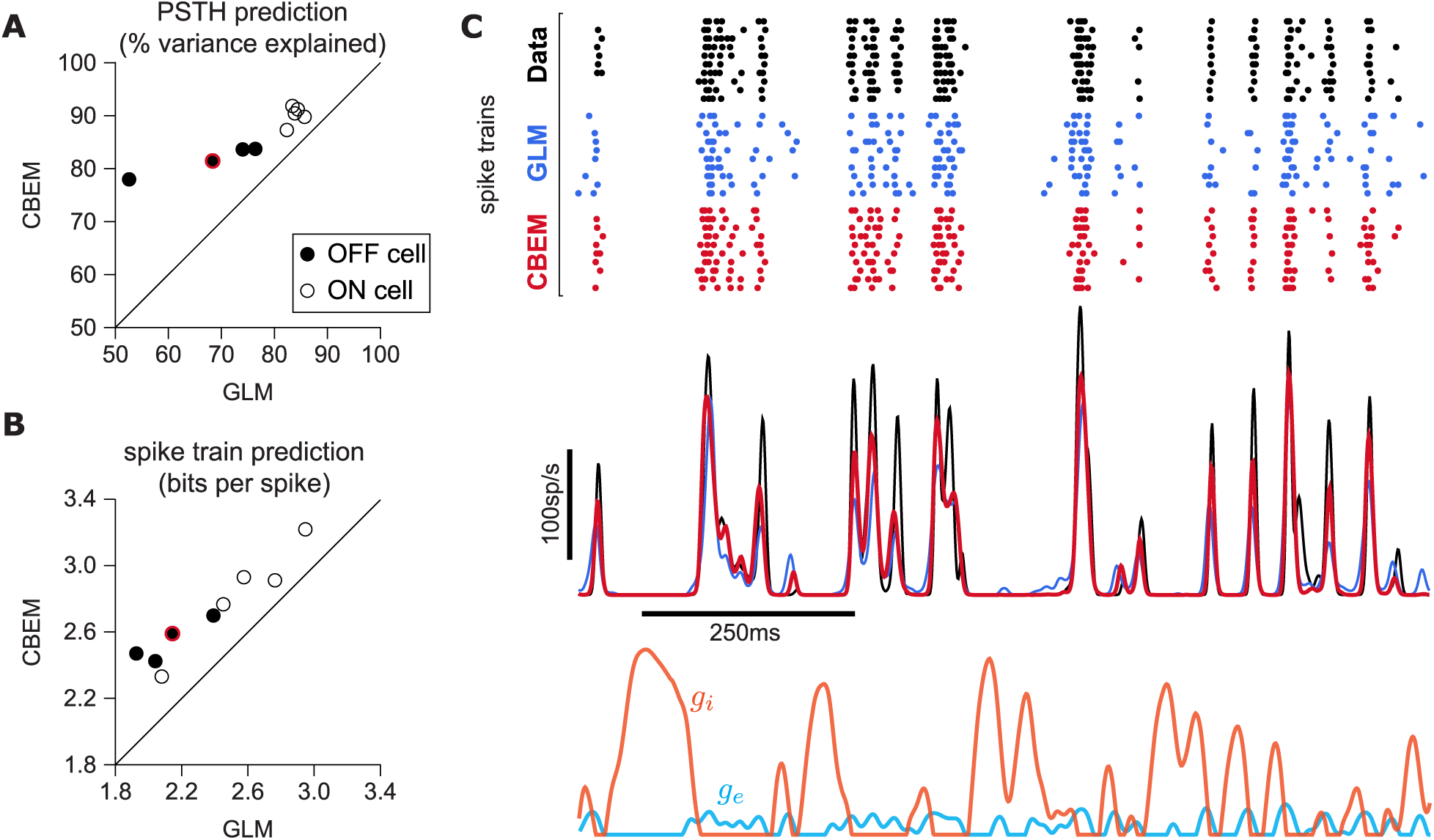
(**A**) Spike rate prediction performance for the population of 9 cells for 5 s test stimulus. The true rate (black) was estimated using 167 repeat trials. The red circle indicates the cell shown in C. (**B**) Log-likelihood of the CBEM compared to the GLM computed on a 5 min test stimulus. (**C**) (top) Raster of responses of an example OFF parasol RGC to repeats of a novel stimulus (black) and simulated responses from the GLM (blue) and the CBEM (red). (middle) Spike rate (PSTH) of the RGC and the GLM (blue) and CBEM (red). The PSTHs were smoothed with a Gaussian kernel with a 2 ms standard deviation. (bottom) The CBEM predicted excitatory (blue) and inhibitory (orange) conductances. The conductances are given in arbitrary units because the model does not include membrane capacitance.

To gain insight into the CBEM’s superior performance, we examined the average firing rate predictions of the GLM along with the average conductance predictions of the CBEM (***Fig. 6c***). We found that GLM rate prediction errors (relative to the PSTH of the real neuron) were anti-correlated with the magnitude of the CBEM inhibitory conductance; the CBEM inhibitory conductance at times when the GLM spike rate exceeded the true spike rate was significantly higher than the CBEM inhibitory conductance at times when the GLM spike rate underestimated the true spike rate (*t*-test, *p <* 0.0001; ***Fig. 6—figure supplement 1b***). This suggests that the CBEM inhibitory conductance helped CBEM predictions by reducing the firing at times when the GLM over-predicted the firing rate. In contrast, the distribution of excitatory conductances did not depend on the sign of the rate prediction error (*t*-test, *p* = 0.19; ***Fig. 6—figure supplement 1a***), and the predicted excitatory conductance was positively correlated with the magnitude of the error (*r* = 0.33, *p <* 0.0001).

Previous experiments have indicated that inhibition only weakly modulates parasol cell responses to full-field Gaussian noise stimuli (Cafaro & Rieke, 2013). To test the effect of inhibition in the model, we also refit the CBEM without any inhibitory synaptic input (CBEM_exc_). We compared the excitatory filters estimated by the CBEM_exc_ with the GLM filters and found that the filters are nearly identical (***Fig. 7e***). This indicates that the GLM stimulus filter accounts only for the excitatory input received by the cell. The CBEM_exc_ still provided a superior prediction of the PSTH than the GLM (81% of the variance explained) and an increased cross-validated log-likelihood (mean improvement of 0.14 ± 0.10bits/sp over the GLM; ***Fig. 7***). The CBEM_exc_ can exhibit changes in total conductance (so it is not technically a GLM, as discussed in Sec 3), and it predicts RGC responses better than the GLM, but not as well as the full CBEM. Thus, the full CBEM achieves superior model performance over the GLM both by including an inhibitory input, and by treating the excitatory input as a conductance-based input in a simple biophysical model.

**Figure 7:**
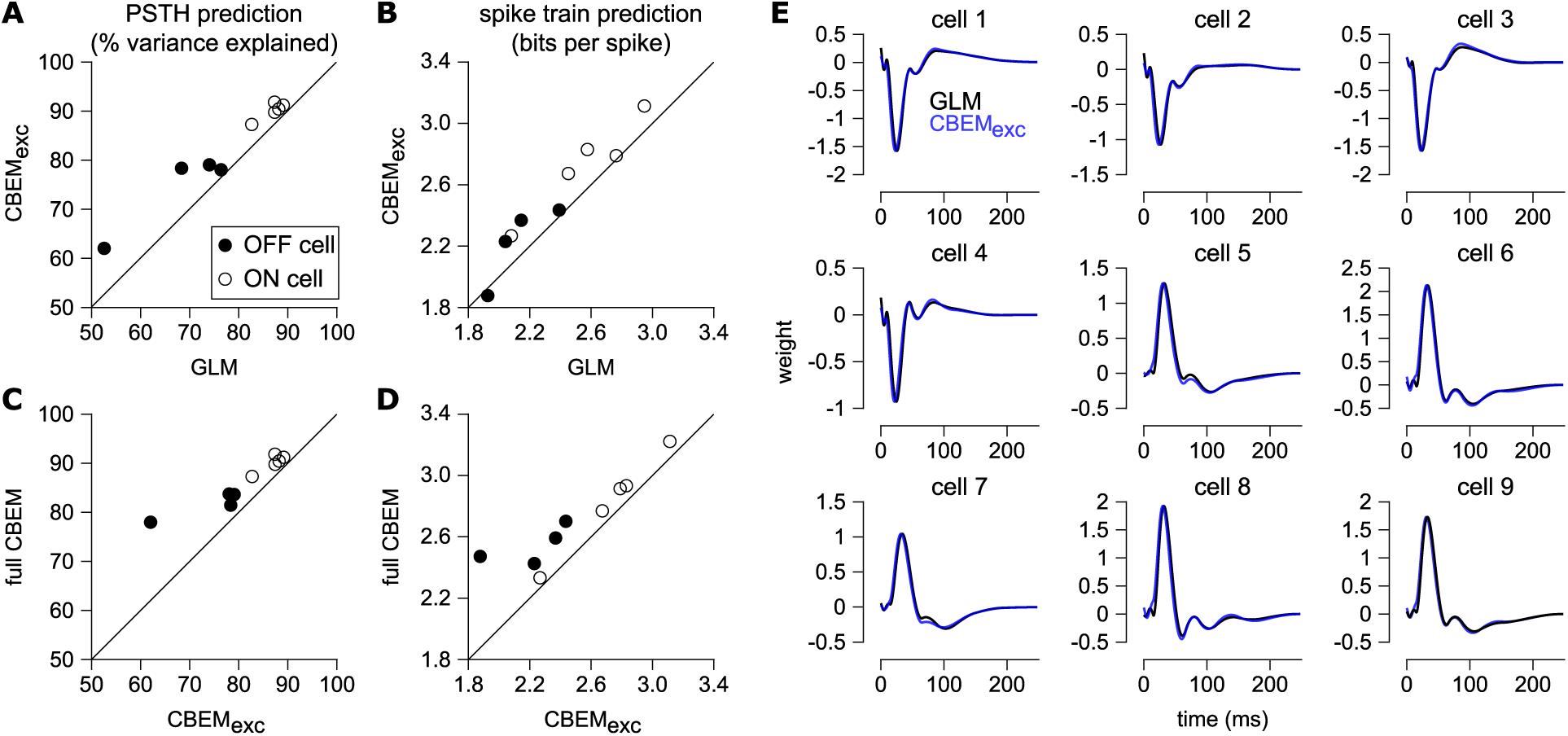
(**A**) Spike rate prediction performance and (**B**) cross-validated log-likelihood for the population of 9 cells for 7 s test stimulus for the GLM and the CBEM with only an excitatory input term (CBEM_exc_). (**C**) The full CBEM with inhibition shows improved spike rate predication and (**D**) cross-validated log-likelihood compared to the model without inhibition. (**E**) The GLM filters for 9 parasol RGCs (black) compared to the excitatory conductance filters estimated by the CBEM without an inhibitory input (blue). The GLM filters are shown scaled to match the height of the CBEM_exc_ filters.

### 7.1 Capturing spike responses across contrasts

Retinal ganglion cells adapt to stimulus statistics such as contrast or variance; increases in stimulus contrast lead to decreases in gain of the neural response, allowing the dynamic range of the response to adapt to the range of contrast values present in the stimulus (Chander & Chichilnisky, 2001; Fairhall et al., 2001; Baccus & Meister, 2002; Beaudoin et al., 2008; Mante et al., 2005; Garvert & Gollisch, 2013; Marava, 2013; Demb & Singer, 2015). Understanding this phenomenon is critical for understanding how the retina codes natural stimuli, because natural scenes vary widely over contrast in both space and time. However, classic linear-nonlinear models with a single linear component fail to capture such effects. This motivates the need for a biophysically plausible modeling framework that can explain RGC responses across stimulus conditions (Ozuysal & Baccus, 2012a; Clark et al., 2013; Cui et al., 2016a).

Previous work has shown that changes in the balance of excitatory and inhibitory input can give rise to multiplicative gain changes in neural responses (Chance et al., 2002; Murphy & Miller, 2003). This raises the possibility that the CBEM may be able account for contrast-dependent changes in RGC responses with single set of parameters. To test this hypothesis, we fit both the CBEM and GLM to eight RGCs stimulated with full-field binary stimuli of 12%, 24%, and 48% contrast. We compared models fit simultaneously to all contrasts with models fit separately to data from each contrast.

To quantify the CBEM’s ability to capture contrast-dependent gain changes in RGC responses, we compared GLM filters fit to RGC responses at each contrast with GLM filters fit to data simulated from the all-contrasts CBEM. Fig. (***Fig. 8a***) shows GLM filters obtained at each contrast for an example RGC, while Fig. ***Fig. 8b*** shows comparable filters fit to spikes simulated from the CBEM fit to this neuron. Both sets of filters exhibit large reductions in amplitude with increasing contrast, the key signature of contrast gain adaptation. Across all eight RGCs, we found high correlation in the filter amplitude scaling for real RGC and simulated CBEM responses (*r* = 0.61, *p <* 0.05; ***Fig. 8—figure supplement 1***).

**Figure 8:**
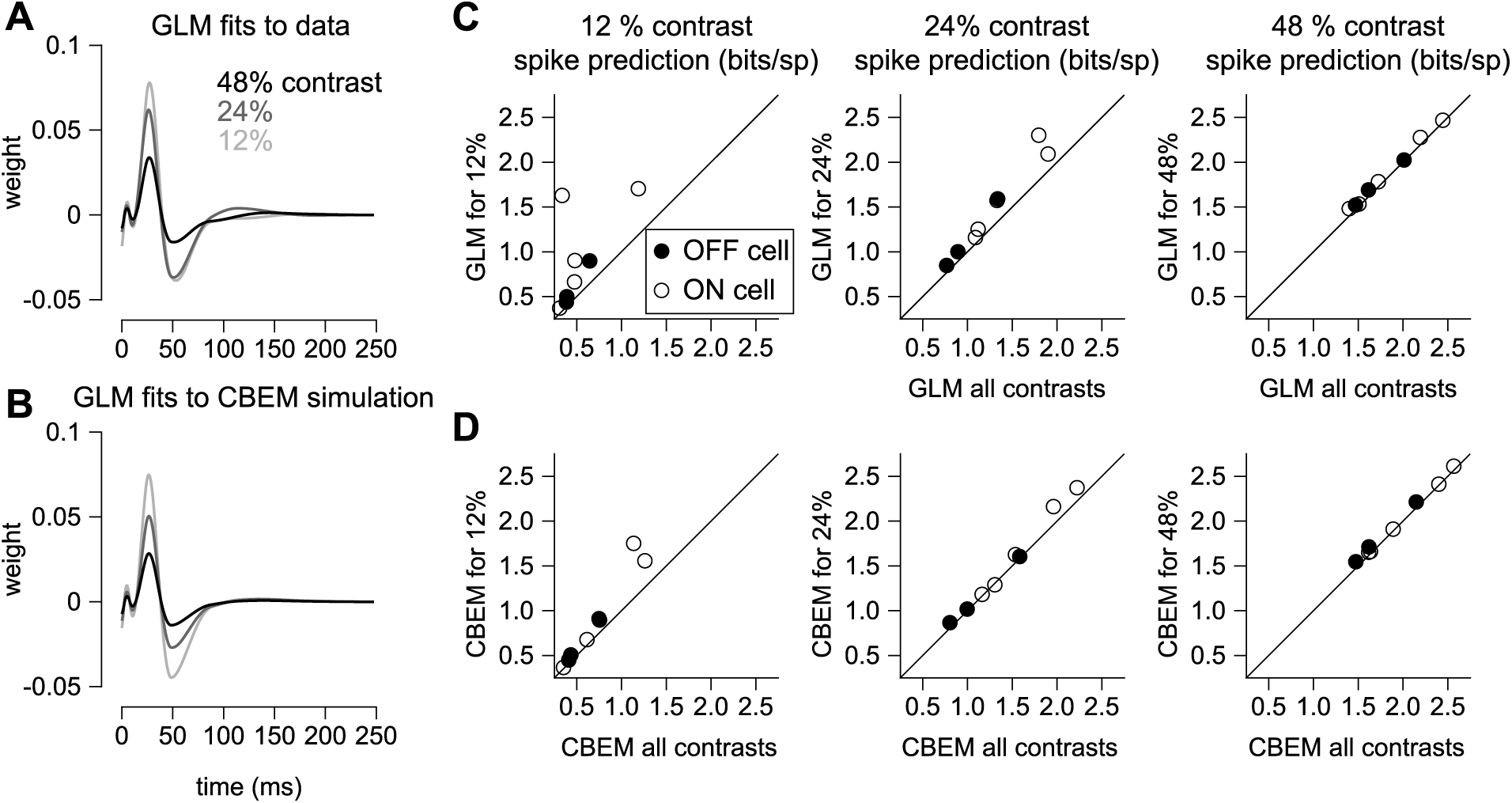
(**A**) GLM filters for an example ON cell fit to responses recorded at 12%, 24%, and 48% contrast. (**B**) GLM filters fit to spike trains simulated from the CBEM fit to the cell shown in A. The CBEM was fit to re-sponses at all 3 contrast levels. Filter height comparisons for CBEM fits to all cells are shown in ***Fig. 8—figure supplement 1.*** Spike train prediction performance of the (**C**) GLM and (**D**) the CBEM tested on a 4 minute stimulus at 12% (left column), 24% (middle column), and 48% (right column) contrast. The model trained on all 3 contrast levels (y-axis) is plotted against the same class of model trained only at the probe contrast level (x-axis).

We found that the CBEM maintained predictive performance across contrast levels more accurately than the GLM (***Fig. 8c-d***). At 12% contrast, the GLM fit to all contrasts lost an average 0.36 ± 0.41 bits/sp (normalized test log-likelihood) compared to GLM fit specifically to the 12% contrast stimulus, while the CBEM lost only 0.16 ± 0.2 bits/sp. At 24% contrast, the GLM lost 0.20 bits/sp while CBEM only lost 0.07 ± 0.14 bits/sp. Finally, both models only lost 0.05 ± 0.08 bits/sp in the 48% contrast probe. The GLM’s partial ability to generalize across these particular conditions despite having only one stimulus filter can be viewed as a consequence of our biophysical interpretation of the GLM; the GLM is equivalent to a biophysical model in which synaptic excitation and inhibition are governed by equal filters of opposite sign; ***Fig. 4 left*** shows that this assumption is approximately correct for ON parasol RGCs. However, the flexibility conferred by the slight differences in these filters gave the CBEM greater accuracy in predicting RGC responses across a range of contrasts.

### 7.2 Capturing spike responses to spatially varying stimuli

To analyze the CBEM’s ability to capture responses to spatially varying stimuli, we examined a dataset of 27 parasol RGCs stimulated with spatio-temporal binary white noise stimuli (Pillow et al., 2008). We fit spatio-temporal filters consisting of a 5−5 pixel field over the same temporal extent as the models fit to full-field stimuli. The temporal profiles of excitatory and inhibitory CBEM filters were qualitatively similar to those that we observed in the full-field stimulus condition (***Fig. 9a,c***). The filters were not constrained to be spatio-temporally separable (the filters were constrained to be rank 2; ***Fig. 9b***), which allowed the synaptic inputs to have different temporal interactions compared to the full-field stimulus.

**Figure 9:**
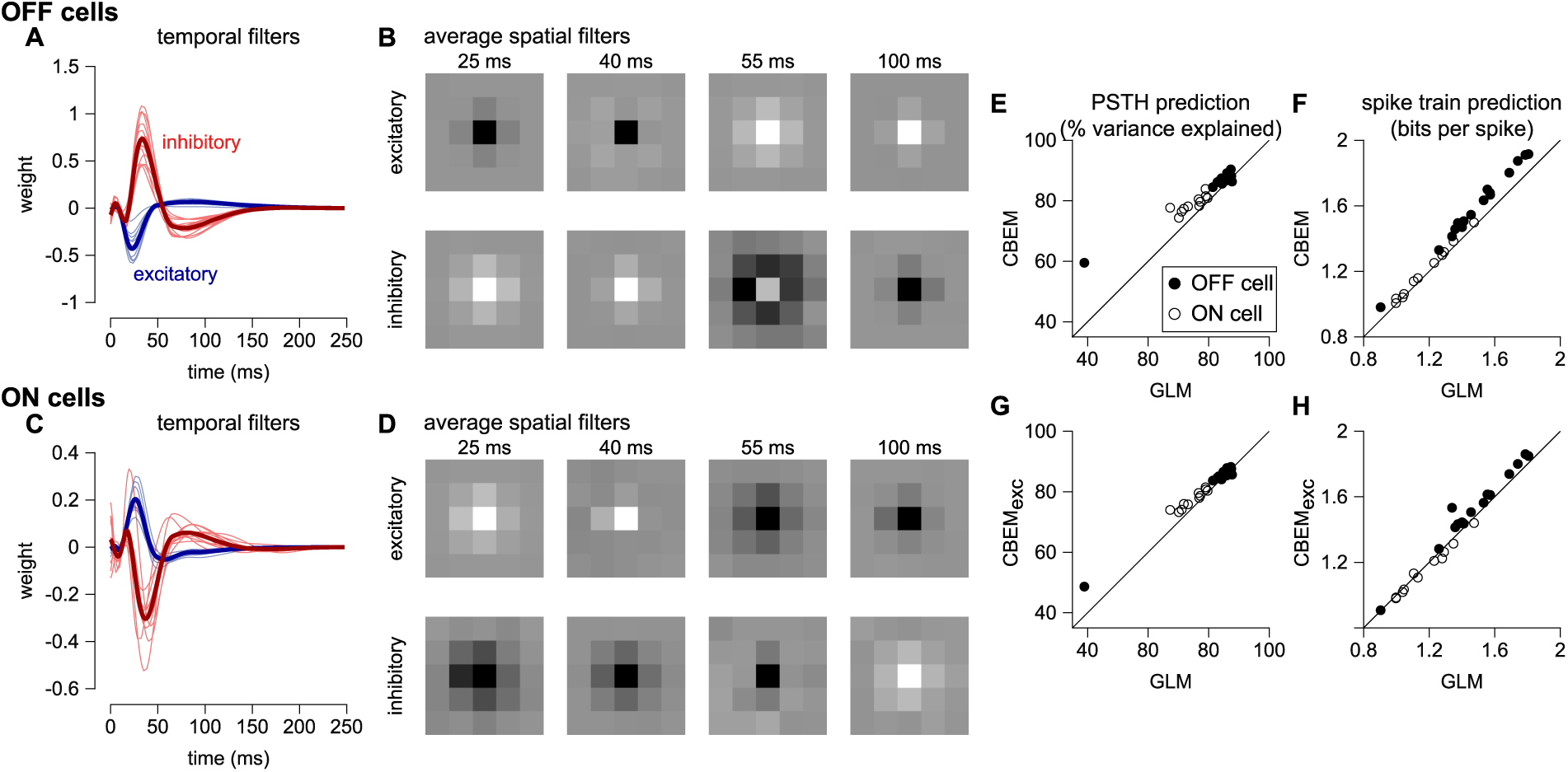
CBEM fits to a population of 27 RGCs. (**A**) Temporal profile of the excitatory (blue) and inhibitory (red) at the center pixel of the receptive field for 16 OFF parasol cells. The thick lines show the mean. (**B**) The mean spatial profiles of the excitatory (top) and inhibitory (bottom) linear filters at four different time points for the OFF parasol cells. (**C,D**) same as A,B for 11 ON parasol cells. (**E**) Spike rate prediction performance of the CBEM compared to the GLM for the population of 27 cells for 8 s test stimulus. The true rate (black) was estimated using 600 repeat trials. (**F**) Log-likelihood of the CBEM compared to the GLM computed on a 5 min test stimulus. (**G**) Spike rate prediction performance of the CBEM_exc_ compared to the GLM. (**H**) Log-likelihood of the CBEM_exc_ compared to the GLM.

We found that the CBEM predicted PSTHs more accurately than a Poisson GLM (83% vs. 79% average R^2^; ***Fig. 9e***). The CBEM also predicted the single-trial responses with higher accuracy than the standard Poisson GLM (average improvement of 0.07±0.04 bits/sp; ***Fig. 9f***). Even the CBEM with excitatory input only yielded more accurate PSTH prediction (81% R^2^) than the GLM, but the single-trial spike train prediction fell to an average of 0.02 ± 0.04 bits/sp higher than the GLM (***Fig. 9g-h***).

To gain insight into how the model’s excitatory and inhibitory inputs shape the CBEM’s responses to spatio-temporal stimuli, we simulated the model with uncorrelated spatio-temporal noise and with spatially correlated stimuli. The uncorrelated spatio-temporal noise was the same independent binary pixel noise used in the RGC recordings, and we used full-field and a binary center-surround stimuli for the spatially correlated noise (***Fig. 10a***). Each frame of the spatially correlated center-surround stimulus was constructed by setting the center pixel to the opposite sign of the pixels in the surround, and the center pixel had equal probability of being black or white. We examined the cross-correlation of the CBEM’s excitatory and inhibitory conductances in each stimulus regime and found that they were similar for the full-field and uncorrelated spatio-temporal noise stimuli (***Fig. 10b*** gray and black traces). In response to these two stimuli, the excitatory and inhibitory conductances showed a strong negative correlation with excitation preceding inhibition (as we saw in ***Fig. 5e***). The center-surround stimulus, however, produced a distinct cross-correlation pattern with a larger positive peak at the positive lags (red traces). Additionally, we simulated GLM and CBEM responses to center-surround contrast steps. The stimulus sequence started as a gray field stepping to a black center pixel with white surround for 500 ms, stepping to a gray field for 500 ms, then stepping to a white center and black surround, finally returning to a gray field (***Fig. 10c*** bottom). The CBEM and GLM showed similar onset responses, but the sustained responses of the CBEM simulations showed inhibition-dependent suppression for both ON and OFF cells (***Fig. 10c*** top and middle). The shape and sustained response of the CBEM_exc_ fit to the OFF cells to center-surround steps qualitatively differed to the full CBEM: the CBEM_exc_ response decayed and then rebounded slightly instead of showing only a decaying response. Thus, full-field and independent spatio-temporal noise result in excitatory and inhibitory correlations that fit closely with the assumptions contained in the GLM. Spatial correlations, and in particular negative correlations, in the stimulus break these assumptions by co-activating excitatory and inhibitory inputs (Cafaro & Rieke, 2013) and therefore spatially correlated stimuli differentiate the CBEM’s predictions from the GLM.

**Figure 10:**
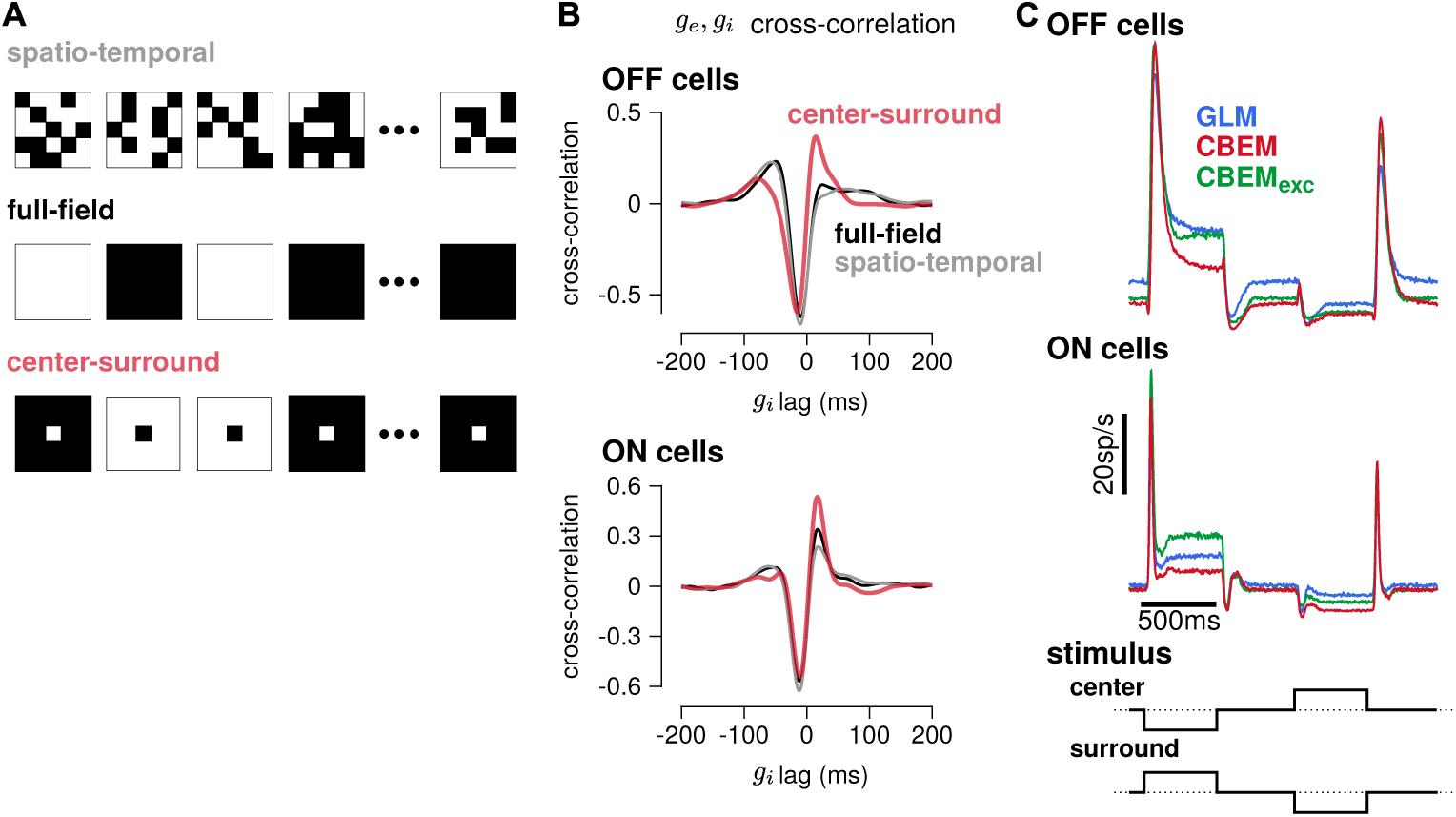
(**A**) Example sequences of 5−5 pixel frames of 3 different types of spatiotemporal noise stimuli used to probe the CBEM. The spatio-temporal stimulus was the same binary noise stimulus used to fit the cells. The full-field stimulus consisted of binary noise at the same contrast and frame rate as the original spatio-temporal stimulus. In the opposing center-surround condition, the center pixel was of opposite contrasts to the surround pixels and the sign of the center pixel was selected randomly on each frame. (**B**) The mean cross-correlation of the CBEM predicted excitatory and inhibitory conductances for the OFF cells (top) and ON cells (bottom) in response to full-field noise (black), spatio-temporal noise (grey), and opposing center-surround noise (red). The strong negative component showed that *g*_*i*_ is delayed and oppositely tuned compared to *g*_*e*_. (**C**) Average firing rate of the GLM (blue), CBEM (red), and CBEM_exc_ (green) fits to 16 OFF cells (top) and 11 ON cells (middle) in response to opposing center-surround contrasts steps (bottom).

## 8 Discussion

The point process GLM has found widespread use for modeling the statistical relationship between stimuli and spike trains. Here we have offered a new biophysical interpretation of this model, showing that it can written as a conductance-based model with oppositely tuned linear excitatory and inhibitory conductances. This motivated us to introduce a more flexible and more biophysically plausible model with independent excitatory and inhibitory conductances, each given by a rectified-linear function of the sensory stimulus. This conductance-based encoding model (CBEM) is no longer technically a general-ized linear model because the membrane potential is a nonlinear function of the stimulus; however, the CBEM has a well behaved point-process likelihood, making it tractable for fitting to extracellular data.

In contrast to purely statistical approaches to designing encoding models, we used intracellular measurements to motivate the choice of the nonlinear functions in the CBEM. We demonstrated that the CBEM not only achieves improved prediction performance compared to the GLM, it accurately recovers the tuning of the excitatory and inhibitory synaptic inputs to RGCs purely from measured spike times. The interaction between excitatory and inhibitory conductances allows the CBEM to change its gain and integration time constant as a function of stimulus statistics (e.g., contrast), an effect that cannot be captured by a standard GLM.

The CBEM belongs to an extended family of neural encoding models that are not technically GLMs because they do not depend on a single linear projection of the stimulus. These include multi-filter LNP models with quadratic terms (Schwartz et al., 2002; Rust et al., 2005; Park & Pillow, 2011; Fitzgerald et al., 2011; Park et al., 2013b; Rajan et al., 2013) or general nonparametric nonlinearities (Sharpee et al., 2004; Williamson et al., 2015); models with input nonlinearities (Ahrens et al., 2008) and multi-linear context effects (Williamson et al., 2016); models inspired by deep learning methods (McIntosh et al., 2016; Maheswaranathan et al., 2017a); and models with biophysically inspired forms of nonlinear response modulation (Butts et al., 2011; Ozuysal & Baccus, 2012b; McFarland et al., 2013; Cui et al., 2016b; Real et al., 2017). The CBEM has most in common with this last group of models, although it stands as the only model so far to have linked spike trains to experimentally measured conductances.

Although the CBEM represents a step towards biophysically realism, it still lacks many biophysical properties of real neurons. For instance, the CBEM’s linear-rectified conductance does not capture the non-monotonic portions of the stimulus-conductance nonlinearities observed in the data (***Fig. 2c,e***); this non-monotonicity likely arises from the fact that amacrine cells can receive inputs from both ON and OFF channels (Manookin et al., 2008; Cafaro & Rieke, 2013). Further developments to the CBEM can include additional sets of nonlinear inputs (McFarland et al., 2013; Maheswaranathan et al., 2017b; Real et al., 2017). Such extensions could include multiple spatially distinct inputs to account for input from different bipolar cells (Freeman et al., 2015; Vintch et al., 2015; Turner & Rieke, 2016; Liu et al., 2017), and spatially selective rectification of inhibitory inputs that helps determine RGC responses to spatial stimuli (Brown & Masland, 2001; Cafaro & Rieke, 2013; Schwartz & Rieke, 2013). Adaptation can occur in localized regions of a ganglion cell’s RF (Garvert & Gollisch, 2013), suggesting that the linear-nonlinear synaptic input functions in the CBEM should be allowed to vary over time. Additionally, future work could apply the CBEM to study the role of active conductances that depend spike history, such as an after hyper-polarization current (Johnston et al., 1995; Badel et al., 2008), and recent work has shown that the parameters of Hodgkin-Huxley style biophysical models can in some instances be recovered from spike trains alone (Meng et al., 2011). Spike-dependent conductances could also be examined in multi-neuron recordings; although the analyses presented here focused on the coding properties of single neurons, many of the RGCs analyzed were recorded simultaneously (Pillow et al., 2008).

Another aspect of the CBEM that departs from biophysical realism is that all stochasticity is confined to the spike generation mechanism. The CBEM models conductances and membrane potential as deterministic functions of the stimulus, which makes the likelihood tractable and allows for efficient fitting with standard conjugate-gradient methods (Real et al., 2017). However, the reliability of RGC spike trains depends on the stochasticity of synaptic conductances (Murphy & Rieke, 2006), and noise correlations between excitatory and inhibitory conductances may also affect encoding in RGCs (Cafaro & Rieke, 2010). A latent variable approach could be used to to incorporate stochasticity in conductances and membrane potential (Meng et al., 2011; Paninski et al., 2012).

Future work will require modeling the neural code using naturalistic stimuli, where the GLM has been shown to fail (Carandini et al., 2005; van Hateren et al., 2002; Butts et al., 2007; Heitman et al., 2016; Turner & Rieke, 2016). Modeling tools must also provide a link between the neural code and computations performed by the neural circuit. As we move towards stimuli with complex spatio-temporal statistics, the ability to connect distinct synaptic conductances to spiking will provide an essential tool for deciphering the complex, nonlinear neural code in sensory systems.

## 9 Materials and methods

### 9.1 Electrophysiology

We analyzed four sets of parasol RGCs. All data were obtained from isolated, peripheral macaque monkey, *Macaca mulatta*, retina.

#### 9.1.1 Synaptic current recordings

We analyzed the responses of 6 ON parasol cells previously described in Trong & Rieke (2008). Cell-attached and voltage clamp recordings were performed to measure spike trains and excitatory and inhibitory currents in the same cells. The stimulus, delivered with an LED, consisted of a one dimensional, full-field white noise signal, filtered with a low pass filter with a 60Hz cutoff frequency, and sampled at a 0.1ms resolution. Spike trains were recorded using 10 unique 6 second stimuli, and each stimulus was repeated 3 or 4 times. After the spike trains were recorded, the excitatory and inhibitory synaptic currents to the same stimuli were measured using voltage clamp recordings. For 4 of the cells, 2-4 trials were recorded for each of the 10 stimuli for the excitatory and inhibitory currents. For the 2 remaining cells, 3-4 excitatory current trials were recorded for all 10 stimuli and 1-2 trials for the inhibitory current were obtained for 8 of the stimuli. Conductances were estimated by dividing the current by the approximate driving force (−70mV for the excitatory currents, and 70mV for the inhibitory).

#### 9.1.2 Dynamic clamp recordings

The membrane potentials of 2 ON parasol retinal ganglion cells were recorded during dynamic clamp experiments previously reported in Cafaro & Rieke (2013). The cells were current clamped and current was injected into the cells according to the equation

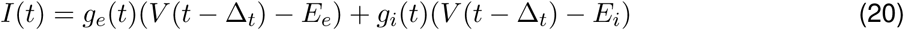

where *g*_*e*_ and *g*_*i*_ were conductances recorded in RGCs in response to a light stimulus. The injected current at time *t* was computed using the previous measured voltage with offset Δ_*t*_ = 100*µ*s. The reversal potentials were *E*_*e*_ = 0 mV and *E*_*i*_ = −90 mV.

For the first cell, 18 repeat trials were recorded for a 19 s stimulation, and 24 repeat trials were obtained from the second cell.

#### 9.1.3 RGC population recordings: full-field stimulus

We analyzed data from two experiments previously reported in Uzzell & Chichilnisky (2004); Pillow et al. (2005). The first experiment included 9 simultaneously recorded parasol RGCs (5 ON and 4 OFF). The stimulus consisted of a full-field binary noise stimulus (independent black and white frames) with a root-mean-square contrast of 96%. The stimulus was displayed on a CRT monitor at a 120Hz refresh rate and the contrast of each frame was drawn independently. A 10 min stimulus was obtained for characterizing the cell responses, and a 5 min segment was used to obtain a cross-validated log-likelihood. Spike rates were obtained by recording 167 repeats of a 7.5 s stimulus.

In a second experiment, 8 cells (5 OFF parasol and 3 on parasol) were recorded in response to a full-field binary noise stimulus (120Hz) at 12%, 24%, and 48% contrast. An 8 min stimulus segment at each contrast level was used for model fitting, and cross-validated log-likelihoods were obtained using a novel 4 min segment at each contrast level.

#### 9.1.4 RGC population recordings: spatio-temporal stimulus

We analyzed 11 ON and 16 OFF parasol RGCs which were previously reported in Pillow et al. (2005). The stimulus consisted of a spatio-temporal binary white noise pattern (i.e., a field of independent white and black pixels). The stimulus was 10 pixels by 10 pixels (pixel size of 120*µ*m − 120*µ*m on the retina), and the contrasts of each pixel was drawn independently on each frame (120Hz refresh rate). The root-mean-square contrast of the stimulus was 96%.

A 10 min stimulus was obtained for characterizing the cell responses, and a 5 min segment was used to obtain a cross-validated log-likelihood. Spike rates were obtained by recording 600 repeats of a 10 s stimulus.

### 9.2 Modeling methods

#### 9.2.1 The conductance-based encoding model

The CBEM introduced above models the spike train response of a RGC to a visual stimulus as a Poisson process where the spike rate is a function of the membrane potential (***Fig. 1b***). The membrane potential is approximated by considering a single-compartment neuron with linear membrane dynamics and conductance-based input (***Eq. 6***). The synaptic inputs (***Eq. 14***) take the form of linear-nonlinear functions of the stimulus, **x**, where *f*_*g*_ is a nonlinear function ensuring positivity of the conductances. The firing rate (***Eq. 19***) is a nonlinear function of the membrane voltage plus a GLM-like spike history (autoregressive) term to account for refractory periods or other spike-dependent behaviors.

The voltage-to-spike rate nonlinearity, *f*_*r*_, follows the form proposed by Mensi et al. (2011), where *µ* is a “soft” spiking threshold and *β* determines the steepness of the nonlinearity. We fixed the values for the firing rate nonlinearity to *α* = 90*sp/s, µ* = −53*mV* and *β* = 1.67*mV*.

Although spiking activity in real neurons influences both the membrane potential and the output nonlinearity (Johnston et al., 1995; Badel et al., 2008), we do not include any spike-dependent currents in the CBEM’s membrane voltage dynamics.

For a set of spike times 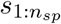 in the interval [0, *S*] and parameters Θ, the log-likelihood in continuous time is

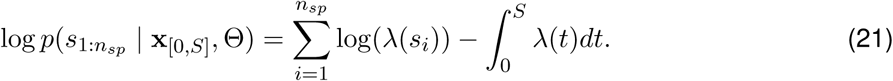

This likelihood can be discretely approximated as the product of *T* Bernoulli trials in bins of width Δ (Citi et al., 2014)

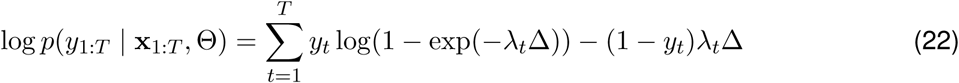

where *y*_*i*_ = 1 if a spike occurred in the *i*th bin and 0 otherwise.

The membrane voltage (and firing rate) is computed by integrating the membrane dynamics equation (***Eq. 6***). In practice, we evaluate *V* along the same discrete lattice of points of width Δ (*t* = 1, 2, 3, *… T*) that we use to discretize the log-likelihood function. Assuming *g*_*e*_ and *g*_*i*_ remain constant within each bin, the voltage equation becomes a simple linear differential equation which we solve according to ***Eq. 16***.

The model parameters we fit were **k**_*e*_, **k**_*i*_, *b*_*e*_, *b*_*i*_, and **h**, which were selected using conjugate-gradient methods to maximize the log-likelihood.

The reversal potential and leak conductance parameters were kept fixed at *E*_*e*_ = 0*mV*, *g*_*l*_ = 200, *E*_*l*_ = −60*mV*, and *E*_*i*_ = −80*mV*. For modeling the cells in which we had access to intracellular recordings, we set the time bin width to Δ = 0.1*ms* to match the sampling frequency of the synaptic current recordings. For the remaining cells, which were recorded in separate experiments, we set Δ = 0.083*ms*, 100 times the frame rate of the visual stimulus.

The stimulus filters spanned over 100 ms, or over 1000 time bins. Therefore, we restricted the excitation and inhibitory filters to a low dimensional basis to limit the total number of free parameters in the model. The basis consisted of 10 raised cosine ‘bumps’ (Pillow et al., 2005, 2008) of the form

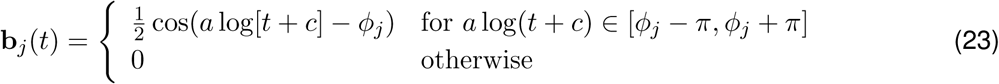

where *t* is in seconds. We set *c* = 0.02 and *a* = *ϕ*_2_ *− ϕ*_1_. The *ϕ*_*j*_ were evenly spaced from *ϕ*_1_ = log(0.0 + *c*), *ϕ*_1_0 = log(0.150 + *c*) so that the peaks of the filters spanned 0*ms* to 150*ms*. The spike history filter was also represented in a low-dimensional basis. The refractory period was accounted for with 5 square basis functions of width 0.4*ms*, spanning the period 0 − 2ms after a spike. The remaining spike history filter consisted of 7 raised cosine basis functions (*c* = 0.0001) with filter peaks spaced from 2*ms* to 90*ms*.

The log-likelihood function for this model is not concave in the model parameters, which increases the importance of selecting a good initialization point compared to the GLM. We initialized the parameters by fitting a simplified model which had only one conductance with a linear stimulus dependence, *g*_*lin*_(*t*) = **k**_*lin*_^*T*^**x**_*t*_ (note that this allowed for negative conductance values). We initialized this filter to 0, and then numerically maximized the log-likelihood for **k**_*lin*_. We then initialized the parameters for the complete model using **k**_*e*_ = *c***k**_*lin*_ and **k**_*i*_ = *−c***k**_*lin*_, where 0 < *c* ≤ 1, thereby exploiting a mapping between the GLM and the CBEM (see Results).

When fitting the model to real spike trains, one conductance filter would occasionally become dominant early in the optimization process. This was likely due to the limited amount of data available for fitting, especially for the cells that were recorded intracellularly. The intracellular recordings clearly indicated that the cells received similarly scaled excitatory and inhibitory inputs. To alleviate this problem, we added a penalty term, *φ*, to the log-likelihood to the *L*_2_ norms of **k**_*e*_ and **k**_*i*_:

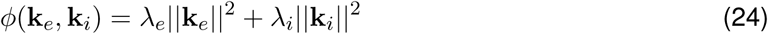

Thus we maximized

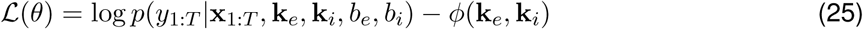

All cells were fit using the same penalty weights: *λ*_*e*_ = 1 and *λ*_*i*_ = 0.2. We note that unlike the typical situation with cascade models that contain multiple filters, intracellular recordings can directly measure synaptic currents. Future work with this model could include more informative, data-driven priors on **k**_*e*_ and **k**_*i*_.

In several analyses, we fit the CBEM without the inhibitory conductance, labeled as the CBEM_exc_. All the fixed parameters used in the full CBEM were held at the same values in the CBEM_exc_.

#### 9.2.2 Fitting the CBEM to simulated spike trains

To examine the performance of our numerical maximum likelihood estimation of the CBEM, we fit the parameters to simulated spike trains from the model with known parameters (***Fig. 1—figure supple-ment 1***). Our first simulated cell qualitatively mimicked experimental RGC datasets, with input filters selected to reproduce the stimulus tuning of macaque ON parasol RGCs (excitation oppositely tuned and delayed compared to excitation, or “crossover” inhibition). The second simulated cell had similar excitatory tuning, but the inhibitory input had the same tuning as excitation with a short delay. The stimulus consisted of a one dimensional white noise signal, binned at a 0.1ms resolution, and filtered with a low pass filter with a 60Hz cutoff frequency. We validated our maximum likelihood fitting procedure by examining error in the fitted filters, and evaluating the log-likelihood on a 5-minute test set. With increasing amounts of training data, the parameter estimates converged to the true parameters for both simulated cells. Therefore, standard fast and non-global optimization algorithms can reliably fit the CBEM to spiking data, despite the fact that the model does not have the concavity guarantees of the standard GLM.

#### 9.2.3 Fitting the conductance nonlinearity

We selected the nonlinear function *f*_*g*_ governing the synaptic conductances by fitting a linear-nonlinear cascade model to intracellularly measured conductances evoked during visual stimulation (Hunter & Korenberg, 1986; Paninski et al., 2012; Park et al., 2013a; Barreiro et al., 2014). We modeled the mean conductance 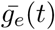 as

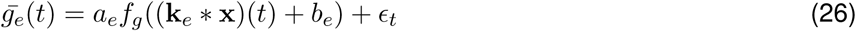

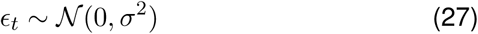

where **x** is a full-field temporal stimulus, and *a*_*e*_ and *b*_*e*_ are constants. We selected a fixed function for the nonlinearity *f*_*g*_. Thus, we chose the **k**_*e*_, *a*_*e*_, and *b*_*e*_ that minimized the squared error between the LN prediction and the measured excitatory conductance.

The soft-rectifying function was selected to model the conductance nonlinearity;

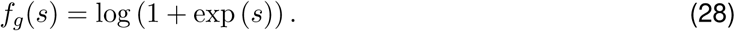

We chose to fix these nonlinearities to known functions rather than fitting with a more flexible empirical form (e.g., Ahrens et al., 2008; McFarland et al., 2013). Fixing these nonlinearities to a simple, closed-form function allowed for fast and robust maximum likelihood parameter estimates while still providing an excellent description of the data.

#### 9.2.4 Fitting the spike-rate nonlinearity

We used a spike-triggered analysis (De Boer & Kuyper, 1968) on membrane voltage recordings to determine the spike rate nonlinearity, *f*_*r*_, as a function of voltage for the CBEM. The membrane potential and spikes were recorded in dynamic-clamp experiments over several repeats of simulated conductances for 2 cells. We computed the mean voltage recorded over all runs of the dynamic-clamp condition, which largely eliminated the action potential shapes from the voltage trace. Using the spike times from all the repeats, we computed the probability of a spike occurring in one time bin given the mean voltage, 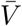:

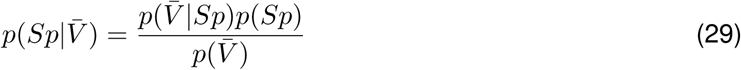

where 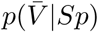 is the spike-triggered distribution of the membrane potential. We combined the spike times and voltage distributions for the two cells to compute a common spike rate function.

A least-squares fit approximated the nonlinearity with a soft-rectification function of the the form

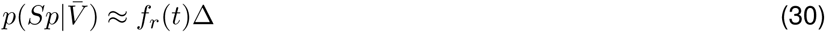

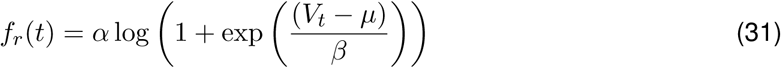

where *α* = 90*sp/s, µ* = −53*mV* and *β* = 1.67*mV*.

We chose to fit the spike-rate nonlinearity with the average voltage recorded over repeat data, instead of looking at the voltage in bins preceding spikes (Jolivet et al., 2006). The average voltage in closer in spirit to the voltage in our model than the single-trial voltage, which contains neither noise nor post-spike currents.

#### 9.2.5 Generalized linear models

For a baseline comparison to the CBEM, we also fit spike trains with a GLM. We used the same Bernoulli discretization of the point-process log-likelihood function for the GLM as we did with the CBEM:

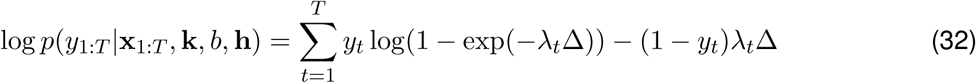

where the firing rate is

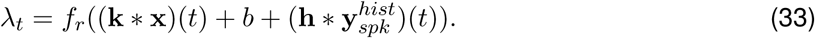

The stimulus filter is **k** and the spike history filter is 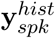 the maximum likelihood estimates for the parameters. We set *f*_*r*_(·) = exp(·), which is the canonical inverse-link function for Poisson GLMs. We found that the soft-rectifying nonlinearity, *f*_*r*_(·) = log(1 + exp(·)), did not capture RGC responses as well as the exponential function (Pillow et al., 2008).

#### 9.2.6 Modeling respones to spatio-temporal stimuli

For spatio-temporal stimuli, the filters for the CBEM and GLM (**k**, **k**_*e*_,, and **k**_*i*_) spanned both space and time. Although the stimulus we used was a 10−10 grid of pixels, the receptive field (RF) of the neurons did not cover the entire grid. We therefore limited the spatial extent of the linear filters to a 5−5 grid of pixels, where the center pixel was the strongest point in the GLM stimulus filter.

The filters were represented as a matrix where the columns span the pixel space and the rows span the temporal dimension. The number of parameters was reduced by decomposing the spatio-temporal filters into a low-rank representation (Pillow et al., 2008). The filter at pixel *x* and time *τ* became

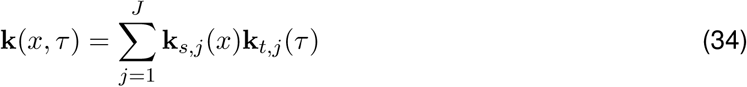

where **k**_*s,j*_ was a vector containing the spatial portion of the filter of length 25 (the number of pixels in the RF) and **k**_*t,j*_ represented the temporal portion of the filter. The temporal filters were projected into the same 10-dimensional basis as the temporal filters used to model the full-field stimuli and the spatial filters were represented in the natural pixel basis. For identifiability, we normalized the spatial filters and forced the sign of the center pixel of the spatial filters to be positive. We used rank 2 filters (*J* = 2) for the CBEM and GLM. Therefore, each filter contained 2 × 25 spatial and 2 × 10 temporal parameters for a total of 70 parameters. In the GLM, we found no significant improvement using rank 3 filters. To fit these low-rank filters, we alternated between optimizing over the spatial and temporal components of the filters.

#### 9.2.7 Evaluating model performance

We evaluated single-trial spike train predictive performance by computing the log-likelihood on a test spike train. We computed the difference between the log (base-2) likelihood under the model and the log-likelihood under a homogeneous rate model (*LL*_*h*_) that captured only the mean spike rate:

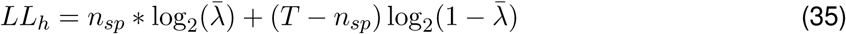

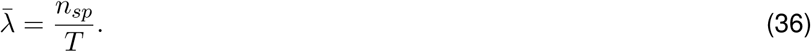

where the test stimulus is of length *T* (in discrete bins) and contains *n*_*sp*_ spikes. We then divided by the number of spikes to obtain the predictive performance in units of bits per spike (bits/sp) (Panzeri et al., 1996; Brenner et al., 2000; Paninski et al., 2004)

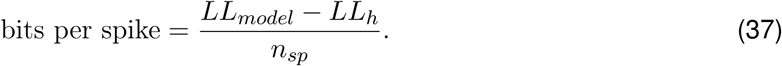

We evaluated model predictions of spike rate by simulating 2500 trials from the model for a repeated stimulus. We computed the firing rate, or PSTH, by averaging the number of spikes observed in 1 ms bins and smoothing with a Gaussian filter with a standard deviation of 2 ms. The percent of variance in the PSTH explained by the model is

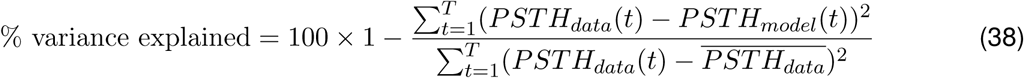

where 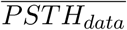 denotes the average value of the PSTH.

## Acknowledgments

We would like to thank EJ Chichilnisky for generously providing data and valuable discussion. We also thank Il Memming Park and Jacob Yates for helpful comments. This work was supported by the McKnight Foundation (JWP), the Simons Foundation (SCGB AWD1004351, JWP), an NSF CAREER Award IIS-1150186 (JWP), a grant from the NIMH (MH099611, JWP), the Howard Hughes Medical Institute (FR), and a grant from the NIH (EY011850, FR).

---

**Figure 1—figure supplement 1:**
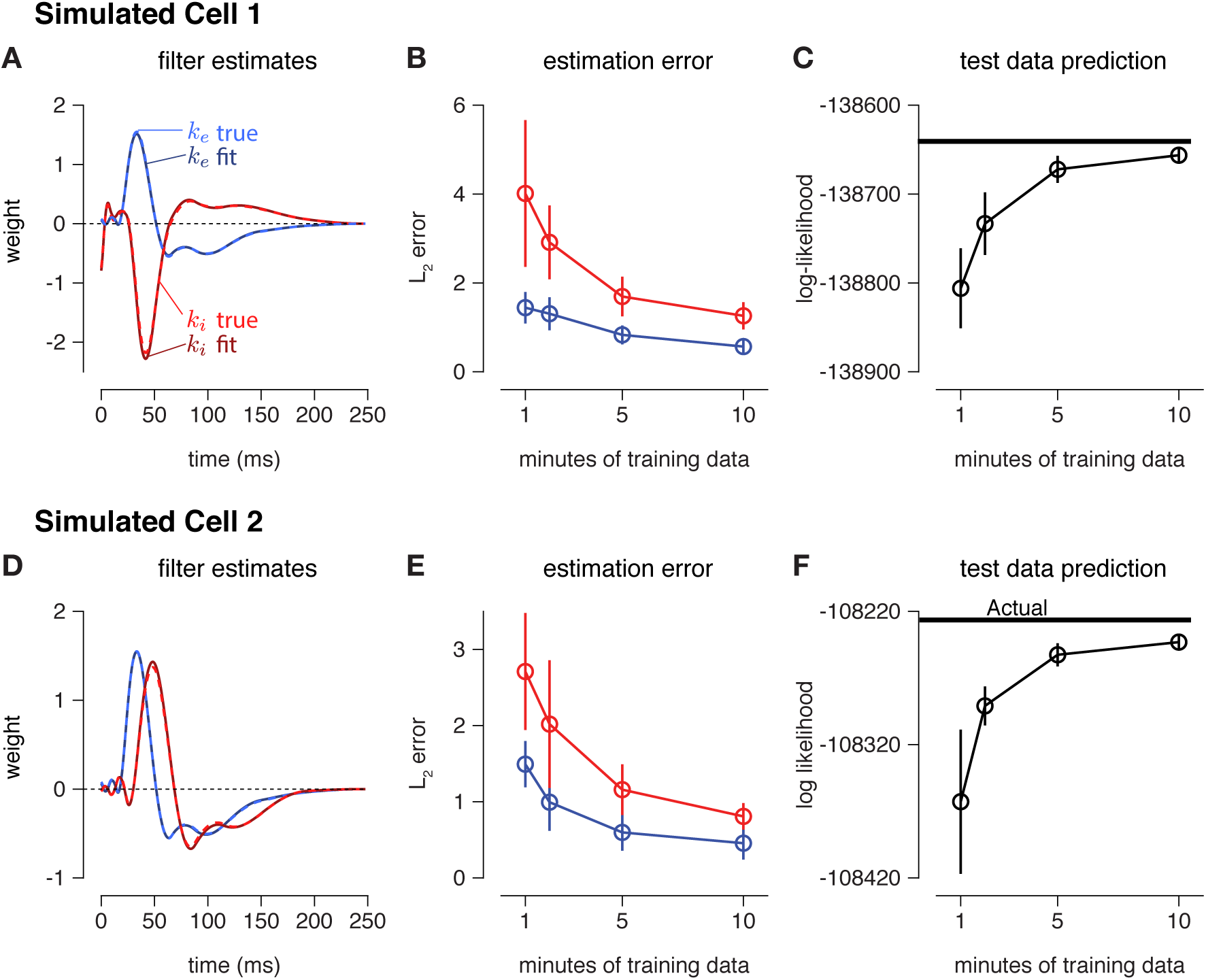
(**A**) Estimates (solid traces) of excitatory (blue) and inhibitory (red) filters from 10 minutes of simulated spike trains. (Dashed lines indicate true filters). The inhibitory filter was oppositely tuned and delayed compared to the excitatory filter. (**B**) The *L*_2_ norm between the estimated input filters and the true filters as a function of the amount of training data. (**C**) The log-likelihood of the fit CBEM on the test data converged towards the log-likelihood of the true model. (**D,E,F**) Same as A-C for a simulation in which **k**_*i*_ had the same tuning as **k**_*e*_ with a delay.

**Figure 6—figure supplement 1:**
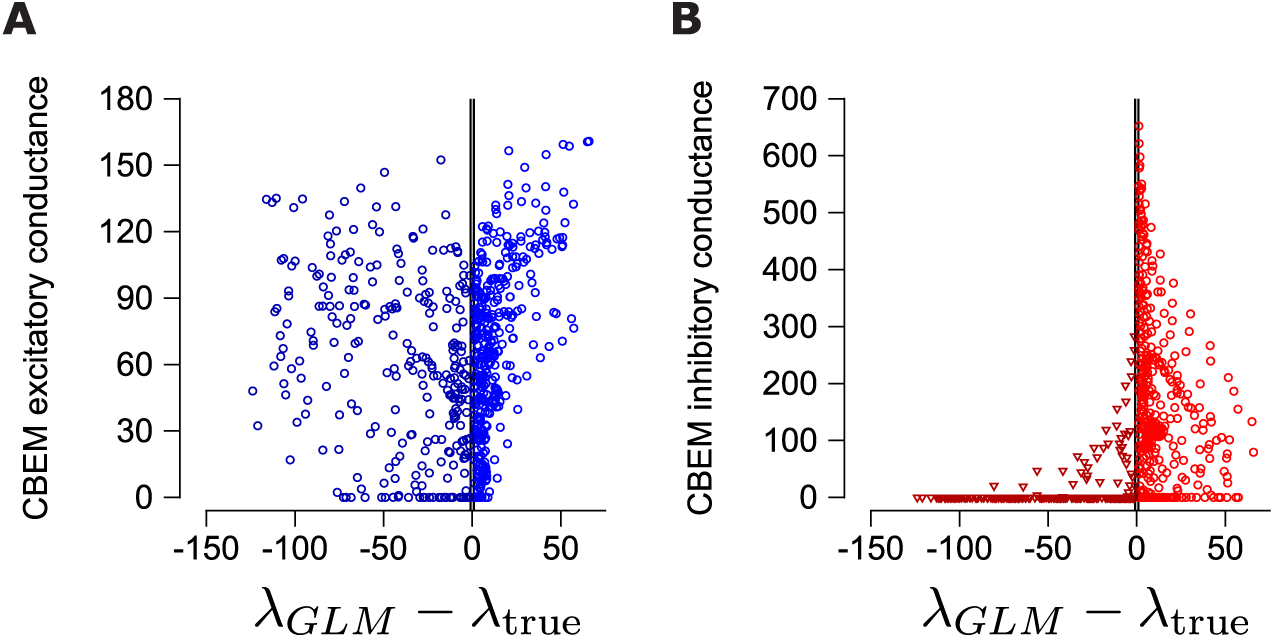
(**A**) The difference in the GLM predicted rate (*λ*_*GLM*_) and the measure spike rate (*λ*_true_) from the PSTH in ***Fig. 6c*** compared to the excitatory conductances predicted by the CBEM. We only considered the time points for which the predicted rate and the measured rate differed by at least 1 spike/sec (gray region denotes the eliminated interval). (**B**) Same as A for the inhibitory conductance.

**Figure 8—figure supplement 1:**
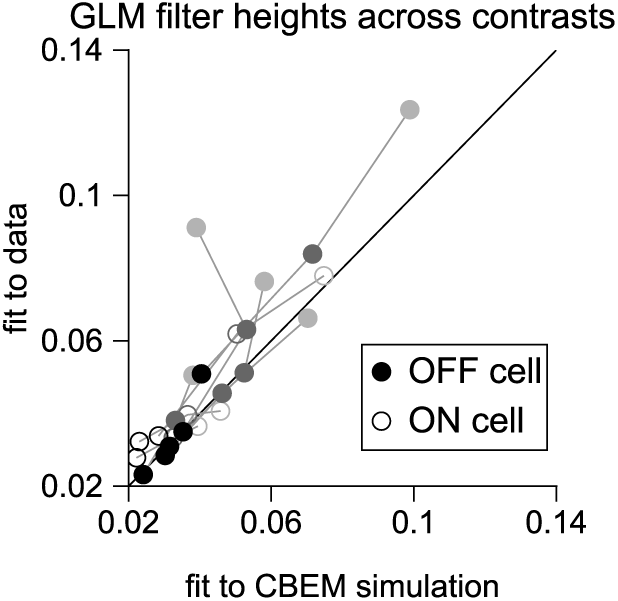
The filter heights (the absolute value of the peak of the filter) of the GLM fits to 8 cells at all three contrast levels (one point per contrast level per cell; lines connect all contrast points from a cell), compared to the GLM filters fit the CBEM simulations of those same cells.

